# A Multi-omic and Multi-Species Analysis of Right Ventricular Failure

**DOI:** 10.1101/2023.02.08.527661

**Authors:** Jenna B. Mendelson, Jacob D. Sternbach, Michelle J. Doyle, Lauren Mills, Lynn M. Hartweck, Walt Tollison, John P. Carney, Matthew T. Lahti, Richard W. Bianco, Rajat Kalra, Felipe Kazmirczak, Charles Hindmarch, Stephen L Archer, Kurt W. Prins, Cindy M. Martin

## Abstract

Right ventricular failure (RVF) is a leading cause of morbidity and mortality in multiple cardiovascular diseases, but there are no approved treatments for RVF as therapeutic targets are not clearly defined. Contemporary transcriptomic/proteomic evaluations of RVF are predominately conducted in small animal studies, and data from large animal models are sparse. Moreover, a comparison of the molecular mediators of RVF across species is lacking. Here, we used transcriptomics and proteomics analyses to define the molecular pathways associated with cardiac MRI-derived values of RV hypertrophy, dilation, and dysfunction in pulmonary artery banded (PAB) piglets. Publicly available data from rat monocrotaline-induced RVF and pulmonary arterial hypertension patients with preserved or impaired RV function were used to compare the three species.

Transcriptomic and proteomic analyses identified multiple pathways that were associated with RV dysfunction and remodeling in PAB pigs. Surprisingly, disruptions in fatty acid oxidation (FAO) and electron transport chain (ETC) proteins were different across the three species. FAO and ETC proteins and transcripts were mostly downregulated in rats, but were predominately upregulated in PAB pigs, which more closely matched the human data. Thus, the pig PAB metabolic molecular signature was more similar to human RVF than rodents. These data suggest there may be divergent molecular responses of RVF across species, and that pigs more accurately recapitulate the metabolic aspects of human RVF.

## Introduction

Right ventricular failure (RVF) is a risk factor for death in multiple cardiovascular diseases including systolic and diastolic heart failure(1), valvular cardiomyopathy(2), congenital heart disease(3,4), and pulmonary hypertension(5,6). Despite these repeated observations, there are currently no RV-specific pharmaceuticals(7), and available therapies for left ventricular (LV) dysfunction are not particularly effective for RVF(7). In addition, the molecular mechanisms underlying RVF are only beginning to be understood and definitively examined, which hampers our ability to define druggable targets(7).

Unquestionably, small animal models are an important tool for dissecting mechanisms of RVF, especially since large animal models require significantly more time and funds. However, the translation of findings obtained from rodents to humans is quite difficult; potentially due to species differences(8), the frequent lack of preclinical rigor(9), and the difficulty performing comprehensive physiological examinations in rodents. Porcine models may offer a more effective translational bridge because the anatomy and physiology of pig hearts are more similar to humans than small animal models(8). Furthermore, advanced imaging and catheterization techniques used in humans are more easily utilized in large animal models(8). To date, most of the molecular evaluations of the pathogenesis of RVF are from small animals, and data from large animals is sparse. Moreover, a comparison of disrupted molecular pathways in rodents, pigs, and humans is lacking.

To address these important knowledge gaps, we implemented a multi-omics approach to define the molecular pathways associated with RV remodeling and dysfunction in a porcine model of RVF induced by pulmonary artery banding (PAB). We integrated RNA-sequencing and proteomics analyses with cardiac MRI derived values of RV size and function in control and PAB male piglets to ascertain which molecular pathways modulate RV remodeling and physiology. Finally, we mined publicly available proteomic and transcriptomic data from the monocrotaline rat model of pulmonary arterial hypertension (PAH)(10), and human PAH patients with compensated and decompensated RV function(11) to perform inter-species comparisons.

## Material and Methods

*Porcine Pulmonary Artery Banding Model*: Four (*n*=4) Yorkshire Cross castrated male piglets aged 4.5±0.6 weeks and weighing 7.3±0.7 kg on the day of surgery were used in this study. Four (*n*=4) pigs of similar age and weight were housed at the University of Minnesota Research Animal Resources facility for 8 weeks and these animals served as controls. Animals were anesthetized and underwent pulmonary artery banding (PAB) surgical procedure using an approach previously described(12). The main pulmonary artery was accessed via a left thoracotomy and an umbilical tape looped around the vessel and cinched until the peak systolic pressure in the right ventricle increased to two-thirds of peak systemic pressure, or approximately 30-40 mmHg. After the ligature was secured, the chest was closed using standard techniques and the animals weaned from anesthesia, recovered and transferred to postoperative housing. In the postoperative period, animals were closely monitored by veterinary staff for clinical signs of right ventricular failure including dyspnea, inappetence, lethargy and ascites on palpation.

*Terminal Data Collection:* After 8 weeks, animals were anesthetized to undergo a terminal cardiac MRI study. Upon completion of the scan, animals were humanely euthanized for a comprehensive gross necropsy and tissue collection. This study was approved by the University of Minnesota Institutional Animal Care and Use Committee (IACUC).

*Cardiac MRI*: We previously described our large animal cardiac MRI protocol(12). Briefly, all studies were performed on a Siemens Aera 1.5T (Siemens, Malvern, PA, USA) scanner. Each examination employed localizers to identify cardiac position and steady-state free procession cine imaging for evaluation of biventricular volumes and function. Segmented short-axis cine images were acquired from the level of the mitral valve to the apex of the left ventricle using a slice thickness of 6 mm. Long-axis cine images were obtained in the four-chamber and three-chamber views. Velocity-encoded imaging was done at the level of the aortic valve and pulmonic valve to confirm ventricular volumes. After cine imaging was completed, contrast-enhanced magnetic resonance angiography evaluated the pulmonary arterial tree. To do this, a small contrast bolus was delivered and then automatic contrast bolus detection was performed to ensure opacification of the main pulmonary artery and the proximal branch pulmonary arteries. The entire examination was gated with electrocardiography. The imaging employed a typical repetition time of 3.0-3.5 ms, echo time of 1.2-1.5 ms, temporal resolution of 35-40 ms, and in-plane spatial resolution of 1.8 x 1.4 mm.

*RNA Isolation*: RNA was isolated with a PureLink RNA Mini kit (Thermo Fisher) with DNase (Zymo Research, Irvine, CA) according to manufacturer’s instructions. Frozen RV tissue from PAB and control piglets was pulverized and incubated in TRIzol^TM^ Reagent (Thermo Fisher) for 5 min. Chloroform was added, and the samples incubated for 3 min. Samples were centrifuged for 15 min at 12,000*g*. The aqueous phase was transferred to a new tube, and an equal volume of 70% ethanol was added. Then samples were centrifuged through the spin cartridge at 12,000*g* for 15 sec. Samples were washed with one half volume of wash buffer I, then 10 µL of DNase I in reaction buffer was added to the column. After 15 min, the column was washed with one half volume of wash buffer I, and twice with wash buffer II. The samples were centrifuged at 12,000*g* for 2 min to dry the membrane. Then 20 μl RNase-free water was added to the center of the spin cartridge, and the cartridge was incubated at 65℃ for 1 minute. Samples were centrifuged at 12,000*g* for 2 min. RNA concentration was determined by spectrophotometry with a NanoDrop 2000(Thermo Fisher), and samples used for bulk RNA-sequencing.

*Bulk RNA-sequencing*: RNA library preparation and sequencing was performed at the University of Minnesota Genomics Center using an Illumina NovaSeq 6000 with 20 million reads per sample.

*Right Ventricular Mitochondrial Enrichments*: RV mitochondrial enrichments were isolated from the same PAB and control piglets with a mitochondrial isolation kit (Abcam, Cambridge, MA) as previously described(10) for TMT 16-plex proteomics analysis.

*Quantitative Mass Spectrometry*: Quantitative mass spectrometry was performed as previously described(10,13) at the University of Minnesota Center for Metabolomics and Proteomics. Mitochondrial enrichment pellets were labeled with TMT16plex isobaric label reagent (Thermo Scientific), cleaned with a 3ml Sep-Pak C18 solid phase extraction cartridge (waters Corporation, Milford, MA), and dried in vacuo. Samples were run on the Eclipse LCMS. LC-MS data was acquired for each sample using a Dionex Ultimate 3000 RSLCnano (Thermo Scientific) in tandem with an Orbitrap Eclipse (Thermo Scientific).

*Integrative Bioinformatics Analysis and Statistics:* Raw sequencing data was trimmed using trimmomatic(14) to remove low quality and Illumina specific adapter sequences. The remaining data was aligned to the pig genome Sscrofa11.1 using HISAT2(15) and gene level count data was produced using subread feature counts and the ensemble gene annotation Sus_scrofa.Sscrofa11.1.106.gtf. Unsupervised clustering of all samples using the top 5,000 most variable genes identified pulmonary artery banded sample CDS19 as an outlier and it was removed from downstream transcriptomic analysis. DESeq2(16) was used to identify differentially expressed genes using counts from protein coding genes only. Heatmaps were produced using the R package pheatmap(17), and volcano plots were produced using the R package EnhancedVolcano(18). R version 4.2.1 was used for all analyses.

Global changes in transcriptomics and proteomics data were assessed using hierarchical cluster analysis, Volcano plots, and random forest classification using R studio and Metaboanalyst(19).

Normalized counts from quantitative mass spectrometry proteomics analysis and bulk RNA-sequencing were used for correlation analyses against cardiac MRI derived values of RV size and function (RV ejection fraction, RV mass, and RV end systolic volume) using Metaboanalyst(19). Proteins and transcripts with |*r*| ≥ 0.5 were extracted and separated with *k*-means clustering (*k*=3) using STRING(20). We selected the top 10 pathways in the KEGG and Reactome databases from each cluster. Pathways that appeared in both the proteomic and transcriptomic analyses were selected for further investigation.

To compare the means of two groups, unpaired *t*-test was used if the data were normally distributed as determined by the Shapiro-Wilk test, and variance was equal as determined by F-test. Mann-Whitney test was used if there was unequal variance between the two groups. Statistical significance was defined as *p*<0.05. Volcano blots, random forest classification, and hierarchical cluster analysis were completed with R version 4.2.1 and MetaboAnalyst(19). Statistical analyses and graphing were performed with GraphPad Prism version 9. Data are presented as mean±standard error of the mean. Graphs show the mean and all individual values.

## Results

### Pulmonary artery banding induced RV remodeling and dysfunction in male piglets

Cardiac magnetic resonance imaging (cMRI, **Supplemental Figure 1**) quantified RV anatomy and function in control (*n*=4) and PAB piglets (*n*=4). As compared to controls, PAB piglets had reduced RV ejection fraction (RVEF, **Figure 1A**), elevated RV mass (**Figure 1B**), increased end systolic (ESV, **Figure 1C**) and diastolic volumes (EDV, **Figure 1D**), and suppressed RV-Pulmonary Artery (PA) coupling (estimated by stroke volume/ESV(21), **Figure 1E**). Additionally, PAB piglets had higher levels of natriuretic peptide B (NPPB, **Figure 1F**). Thus, PAB piglets exhibited adverse RV remodeling and significant RV dysfunction.

**Figure 1:**
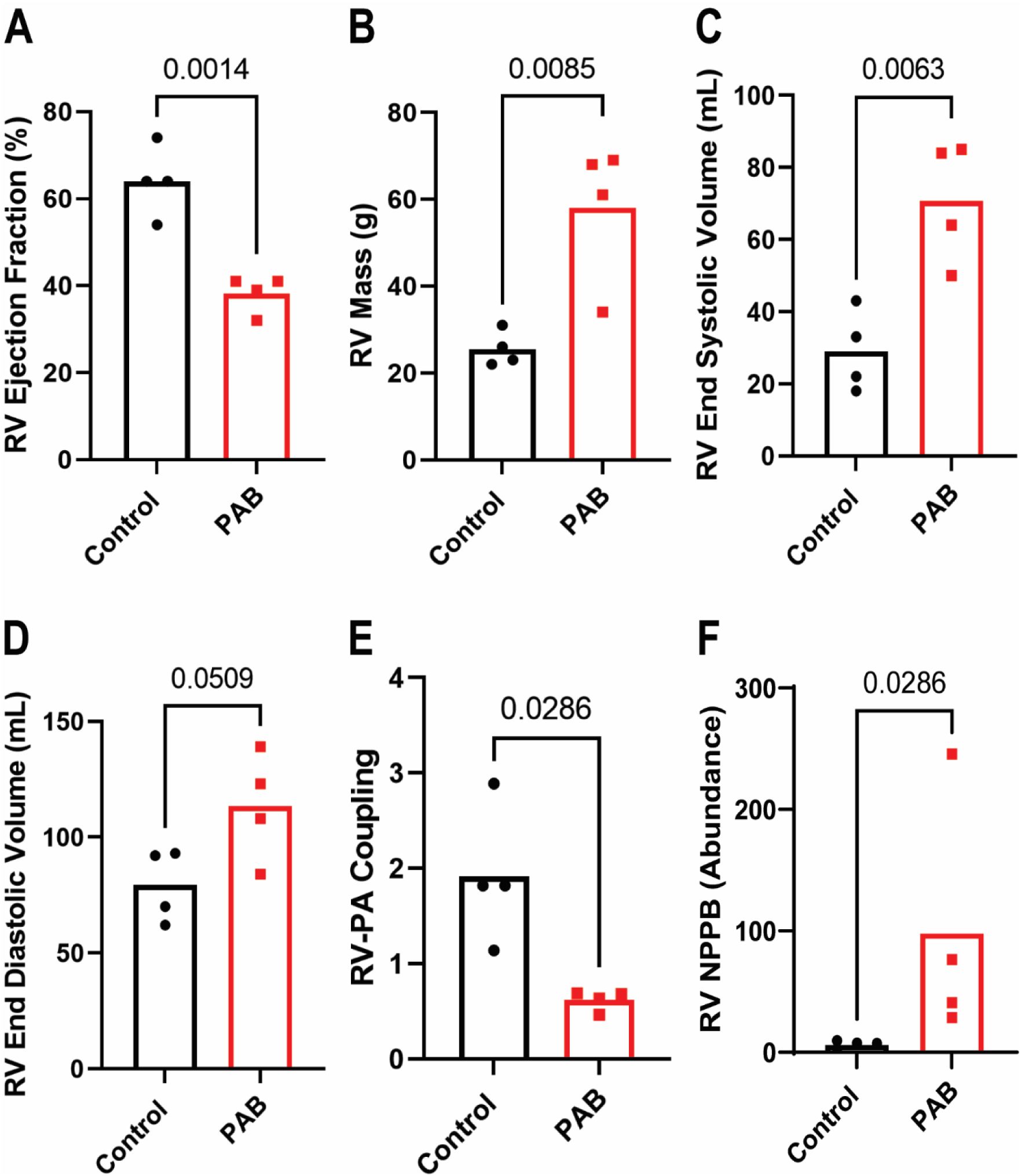
Cardiac MRI Identified Changes in RV Morphology and Function in PAB piglets. PAB piglets have reduced RVEF (control: 64.0±4.1, PAB: 38.3±2.1, *p*=0.014) (**A**), elevated RV mass (control: 25.5±2.0, PAB:58.0±8.2, *p*=0.014) (**B**), RV dilation RV ESV (control: 29.0±5.6, PAB: 70.8±8.4, *p*=0.014) (**C**) and RV EDV control: 79.3±7.8, PAB:113.5±11.7, *p*=0.014) (**D**), impaired RV-PA coupling (control: 1.9±0.4, PAB:0.6±0.1, *p*=0.014) (**E**), and elevated NPPB (control: 6.1±2.1, PAB: 97.9±50.3) (**F**) . *p*-values determined by unpaired t-test (**A-E**), and Mann-Whitney test (**F**) when there was unequal variance between groups.

### Transcriptomics and proteomics analyses identified global changes in the RV of PAB piglets

We next performed a multi-omic analysis using transcriptomics and proteomics to define the molecular signature of porcine RVF. Hierarchical cluster analysis (**Figure 2A**) of transcriptomic data demonstrated the two experimental groups were distinct. Filtering the data using an adjusted *p*-value of 0.05, we identified 1530 transcripts that were significantly different between the two groups with 792 being elevated and 738 being reduced in PAB piglets (**Figure 2B**). Random forest classification identified the following transcripts that differentiated the two experimental groups (**Figure 2C**): SRY-box transcription factor 18 (SOX18), dachshund family transcription factor 1 (DACH1), elongation factor for RNA polymerase II 3 (ELL3), non-specific cytotoxic cell receptor protein 1 (NCCRP1), cathepsin D (CTSD.1), collagen type XCIII alpha 1 chain (COL18A1), cytoplasmic linker associated protein 2 (CLASP2), baculoviral inhibitor of apoptosis repeat containing 5 (BIRC5), BCL6 corepressor like 1 (BCORL1), ankyrin repeat and suppressor of cytokine signaling box containing 12 (ASB12), alsin rho guanine nucleotide exchange factor (ALS2), asparagine-linked glycosylation 14 UDP-N-acetylglucoasminyltransferase subunit (ALG14), activated leukocyte cell adhesion molecule (ALCAM), adrenoreceptor alpha 1A (ADRA1A), and a disintegrin and metalloproteinase domain 22 (ADAM22). 10 out of 15 transcripts (SOX18, DACH1, NCCRP1, CLASP2, BCORL1, ASB12, ALS2, ALCAM, ADRA1A, and ADAM22) were decreased in PAB piglets, and 5 (ELL3, CTSD.1, COL18A1, BIRC5, and ALG14) were increased.

**Figure 2:**
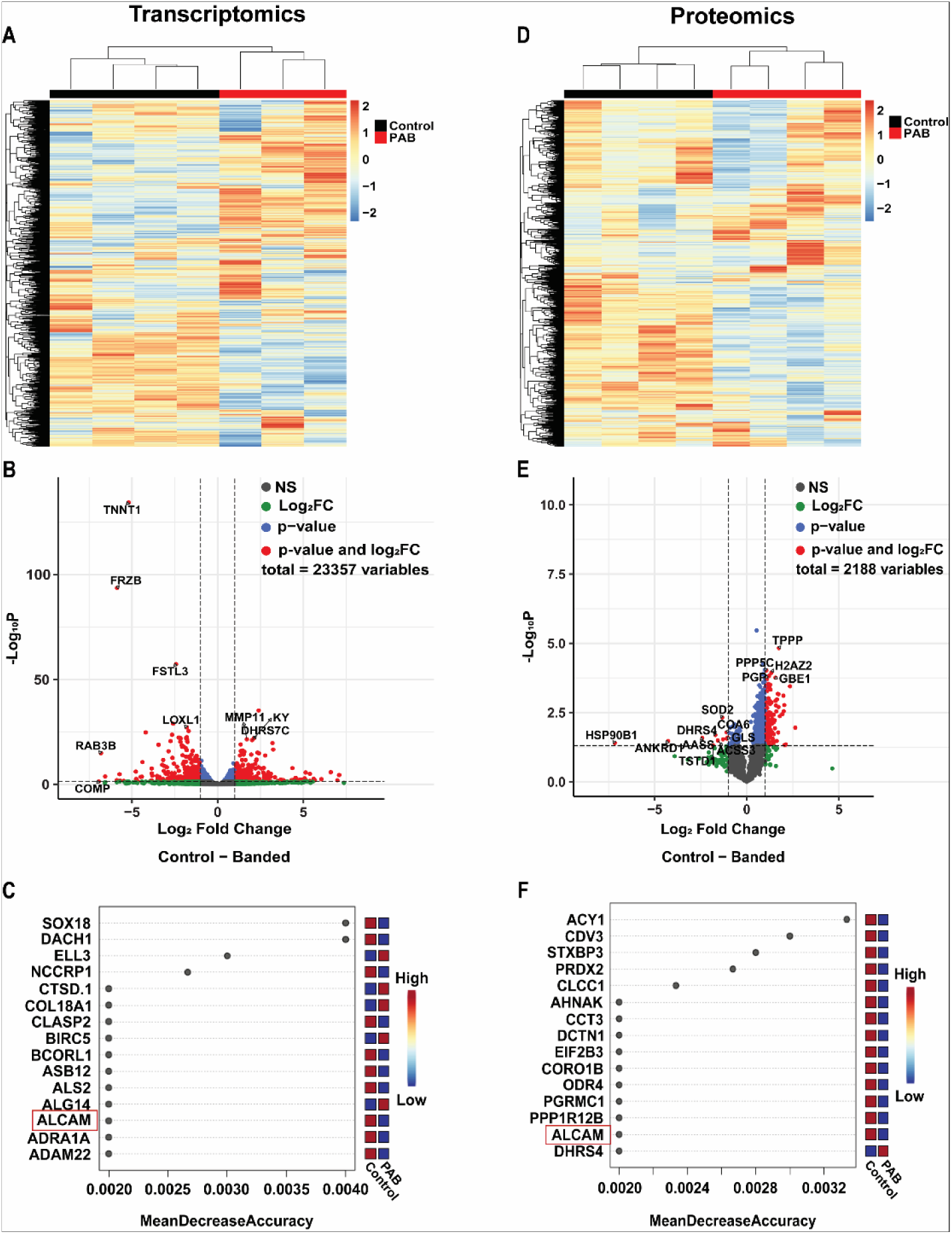
Transcriptomics and Proteomics Analyses Demonstrated PAB RVs Exhibit a Distinct Molecular Signature as Compared to Controls. Hierarchical cluster analysis (**A**) of transcriptomic data reveal control and PAB piglets are molecularly distinct. Volcano plot (**B**) found 1530 transcripts were different between control and PAB piglets. 792 were elevated and 738 were reduced in PAB piglets. The top 15 most important transcripts for distinguishing between control and PAB piglets as determined by Random Forest analysis (**C**). Hierarchical cluster analysis of proteomics data demonstrate distinct profiles when comparing control and PAB piglets (**D**). 710 proteins were upregulated and 103 were downregulated by volcano blot (**E**). The top 15 most important proteins for differentiating between control and PAB piglets as determined by Random Forest analysis (**F**).

Hierarchical cluster analysis of proteomics data revealed distinct profiles when the two groups were compared (**Figure 2D**). 710 proteins were upregulated and 103 proteins were downregulated in PAB RVs (**Figure 2E**). Random forest classification correctly sorted all 4 control samples, and 3 of the 4 PAB samples. The most important proteins for differentiating the two groups (**Figure 2F**) were: aminoacylase 1(ACY1), carnitine deficiency-associated gene expressed in ventricle 3 (CDV3), syntaxin binding protein 3 (STXBP3), peroxiredoxin 2 (PRDX2), chloride channel CLIC like 1 (CLCC1), neuroblast differentiation-associated protein (AHNAK), chaperonin containing TCP1 subunit 3 (CCT3), dynactin subunit 1 (DCTN1), eukaryotic translation initiation factor 2B subunit gamma (EIF2B3), coronin 1B (CORO1B), ODR4 GPCR localization factor homolog (ODR4), progesterone receptor membrane component 1 (PGRMC1), protein phosphatase 1 regulatory subunit 1B (PPP1R12B), activated leukocyte cell adhesion molecule (ALCAM), dehydrogenase/reductase 4 (DHRS4). 14 out of 15 proteins (ACY1, CDV3, STXBP3, PRDX2, CLCC1, AHNAK, CCT3, DCTN1, EIF2B3, CORO1B, ODR4, PGRMC1, PPP1R12B, and ALCAM) that differentiated the groups were downregulated in PAB piglets as only DHRS4 was upregulated in PAB piglets. In both the proteomic and transcriptomic analyses, ALCAM was decreased in PAB piglets and helped differentiate the groups.

### Integration of transcriptomics, proteomics, and cardiac MRI identified pathways related to RV dysfunction, hypertrophy, and dilation

To provide more granularity about the molecular pathways that contribute to adverse RV remodeling and RV dysfunction in our PAB porcine model, we performed Kyoto Encyclopedia of Genes and Genomes (KEGG) and Reactome pathway analyses on transcripts and proteins that were correlated with cMRI-derived evaluation of RV function, hypertrophy, and size (**Supplemental Figures 2-13**). The pathways that were associated with RVEF in both proteomics and transcriptomics approaches were Lysosome, Metabolism of Lipids, and Fatty Acid Metabolism (**Figure 3A-C**). In the Lysosome pathway, most molecules were negatively associated with RVEF. Niemann-Pick C1 protein (NPC1), a lysosomal membrane protein involved in intracellular cholesterol trafficking(22), appeared in both the transcriptomic and proteomic analyses and was negatively correlated with RVEF. In the Metabolism of Lipids and Fatty Acid Metabolism pathways, there were transcripts and proteins that were positively and negatively associated with RVEF(**Figure 3** **B and C**), suggesting these pathways were altered in a bimodal manner.

**Figure 3:**
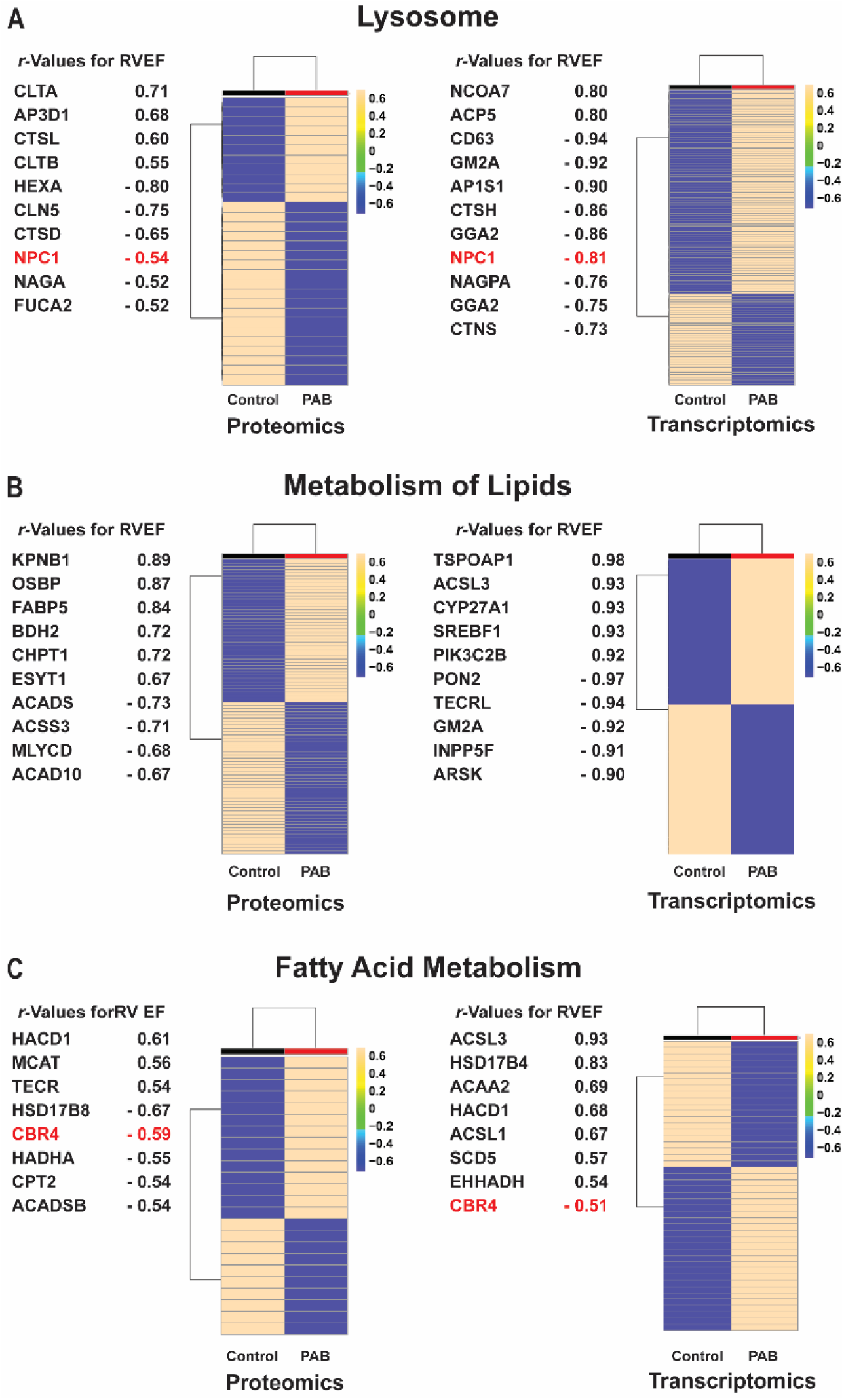
Correlational analysis of proteomics and transcriptomics data identified Lysosome, Metabolism of Lipids, and Fatty Acid Metabolism as key pathways associated with RVEF. Proteins (left) and transcripts (right) *r*-value for RVEF in the (**A**) Lysosome, (**B**) Metabolism of Lipids, and (**C**) Fatty Acid Metabolism pathways. Hierarchical cluster analysis of proteomics and transcriptomics data demonstrate differences between control and PAB piglets. Molecules identified in both proteomics and transcriptomics analyses were highlighted in red.

Next, we found the Dilated Cardiomyopathy and Arrhythmogenic Right Ventricular Cardiomyopathy (ARVC) pathways were associated with RV hypertrophy (**Figure 4A and B**). In the proteomics data, most proteins in both pathways were downregulated in PAB animals, but in the transcriptomics data, there were bidirectional changes. Sarcoplasmic/endoplasmic reticulum calcium ATPase 2 (ATP2A2) and CACNB2, a subunit of L-type voltage gated Ca channels, were identified in both transcriptomic and proteomic analyses. In both experimental methods, ATP2A2 and CACNB2 were downregulated in the setting of RV hypertrophy. Integrin subunit alpha V (ITGAV) was identified in both analyses, but ITGAV was negatively correlated with RV mass in the proteomic analysis, but positively correlated with RV mass in the transcriptomic analysis.

**Figure 4:**
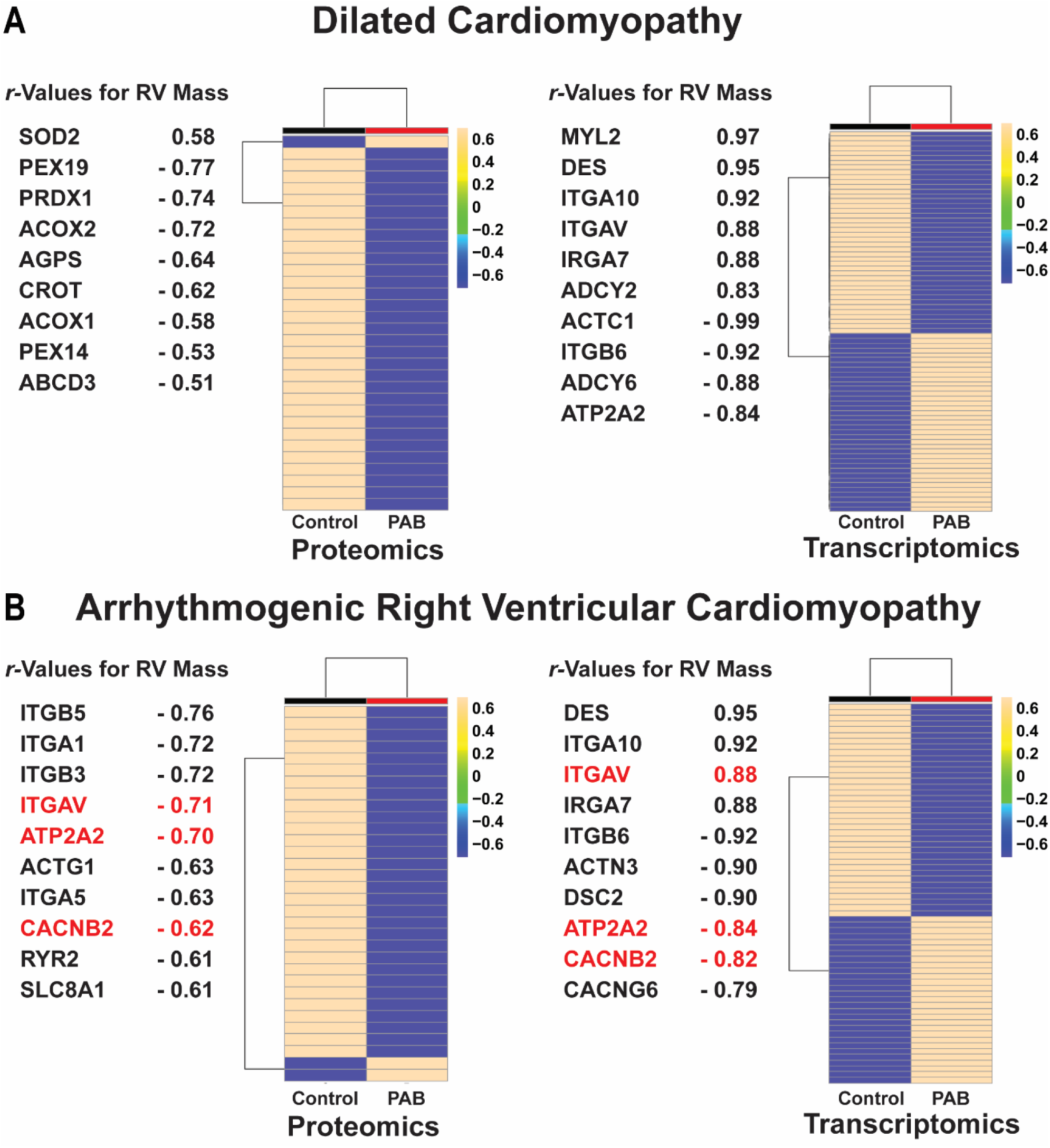
Correlational analysis of proteomics and transcriptomics data identified Dilated Cardiomyopathy and Arrhythmogenic Right Ventricular Cardiomyopathy as key pathways associated with RV hypertrophy. Proteins (left) and transcripts (right) with *r*-value for RV mass in the (**A**) Dilated Cardiomyopathy, and (**B**) Arrhythmogenic Right Ventricular Cardiomyopathy pathways. Hierarchical cluster analysis of proteomics and transcriptomics data demonstrate differences between control and PAB piglets. Molecules identified in both proteomics and transcriptomics analyses were highlighted in red.

When evaluating pathways associated with RV dilation as defined by ESV, we identified Metabolism, Respiratory Electron Transport, and Citric Acid Cycle and Respiratory Electron Transport pathways are predictors of adverse RV remodeling (**Figure 5 A, B, and C**). There were bidirectional changes in molecules in all of these pathways, but RV dilation was associated with higher levels of multiple components of the electron transport chain (ETC). In particular, cytochrome C oxidase subunit 6A1 (COX6A1), synthesis of cytochrome C oxidase 1 (SCO1), dihydrolipoamide S-succinyltransferase (DLST), and NADH-ubiquinone oxidoreductase complex (NDUFV2) proteins were increased in PAB animals in the proteomics-based evaluation. In the transcriptomic analysis, abundances of lactate dehydrogenase-A (LDHA), mitochondrial encoded ATP synthase membrane subunit 6 (ATP6), cytochrome C oxidase I (COX1), mitochondrially encoded NADH:ubiquinone oxidoreductase core subunit 1 (ND1), mitochondrially encoded NADH:ubiquinone oxidoreductase core subunit 3 (ND3), and ubiquinol-cytochrome C reductase complex III subunit VII (UQCRQ) were elevated in the PAB RV.

**Figure 5:**
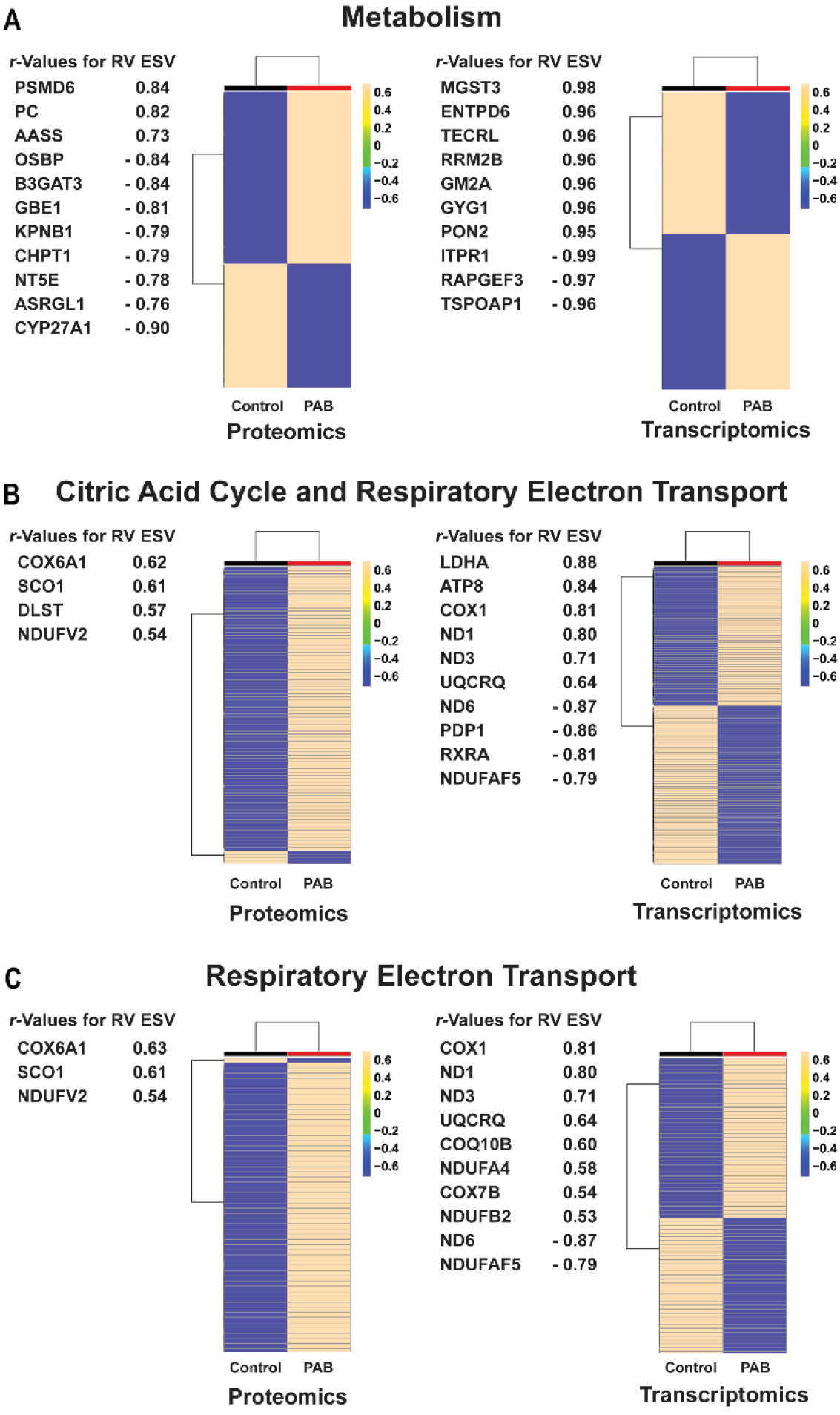
Correlational analysis of proteomics and transcriptomics data identified Metabolism, Citric Acid Cycle and Respiratory Electron Transport, and Respiratory Electron Transport pathways associated with RV ESV. Proteins (left) and transcripts (right) with *r*-value for RV ESV in the (**A**) Metabolism, (**B**) Citric Acid Cycle and Respiratory Electron Transport and (**C**) Respiratory Electron Transport Pathways. Hierarchical cluster analysis of proteomics and transcriptomics data demonstrate differences between control and PAB piglets.

### Multi-species comparisons suggested pigs exhibited a metabolic molecular signature that was more closely related to human RV failure than rodents

Because Fatty Acid Oxidation (FAO) and ETC pathways were identified in our porcine analysis and alterations in these pathways are present in human heart failure(23,24), we evaluated and compared the relative changes in FAO and ETC in rodents (GSE119754, PXD027273), pigs, and humans (GSE198618) (10,11,25). In the proteomics analysis, abundances of nearly all FAO and ETC proteins were reduced in MCT rats as compared to control rats (**Figure 6A-B**). However, FAO and ETC proteins were mostly upregulated in PAB pigs when compared to control pigs. Human patients with compensated RV function (cRV) displayed suppression of many FAO and ETC proteins, but patients with decompensated RV (dRV) function had higher abundances of most proteins. In the transcriptomic analysis, MCT rats again had lower levels of FAO and ETC transcripts compared to control rats. In PAB pigs, FAO and ETC transcripts changed in a bidirectional manner. Patients with cRV had decreased abundances of most FAO and ETC transcripts, but patients with dRV predominately displayed increased abundances of transcripts in these two pathways. Thus, these data suggest PAB pigs have a metabolic molecular signature that more closely mimics human RVF.

**Figure 6:**
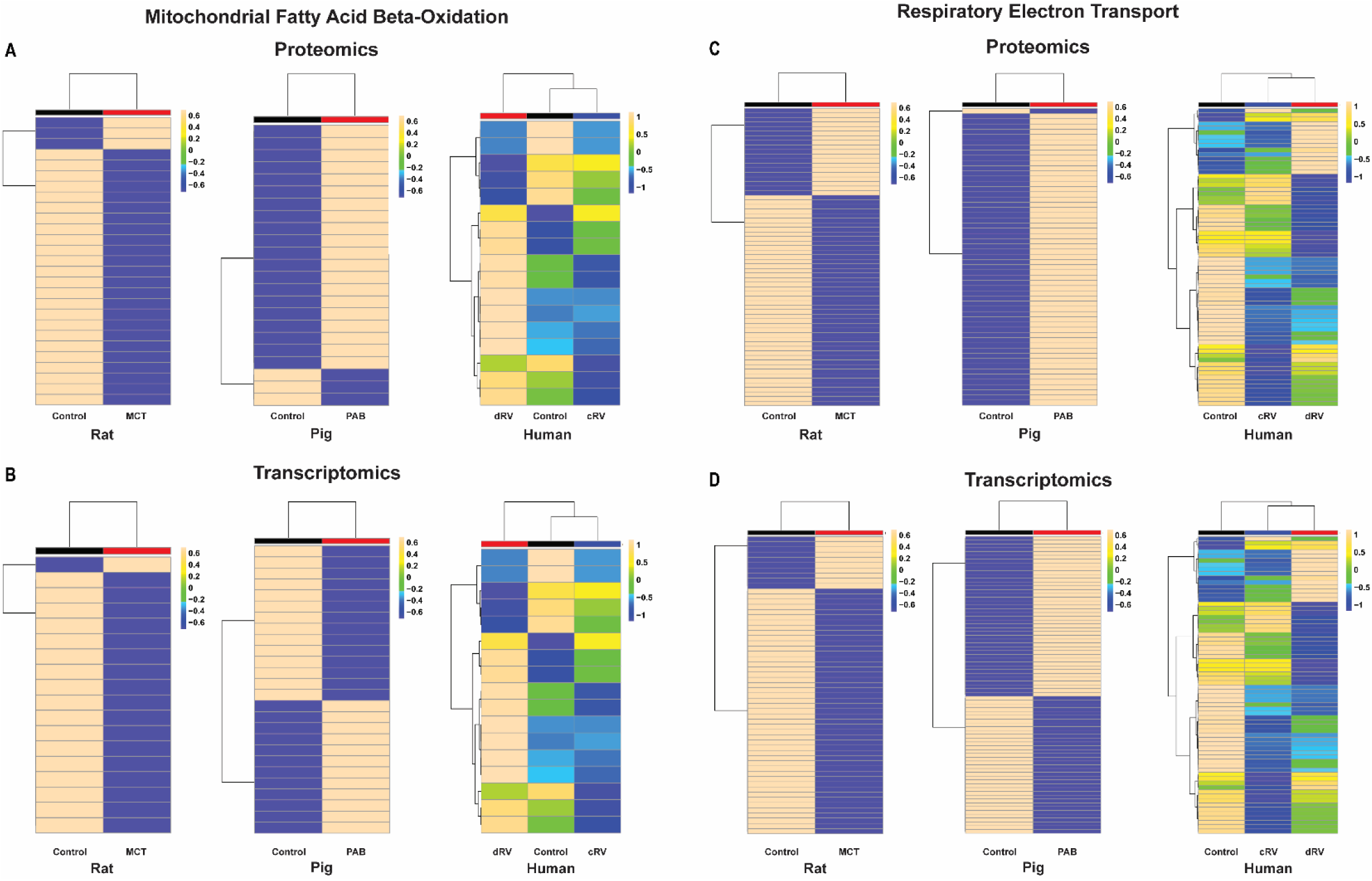
Comparison of Mitochondrial Fatty Acid Beta-Oxidation and Respiratory Electron Transport Pathways Across Species. Hierarchical cluster analyses of proteomics data from rat, pig, and human experiments analyzing the proteins in the (**A**) Mitochondrial Fatty Acid Beta-Oxidation and (**C**) Respiratory Electron Transport pathways. Hierarchical cluster analyses of transcriptomics data from rat, pig, and human experiments analyzing the (**B**) Mitochondrial Fatty Acid Beta-Oxidation and (**D**) Respiratory Electron Transport pathways.

## Discussion

In this large animal study, cMRI demonstrates PAB induces significant RV hypertrophy, dilation, and dysfunction. The adverse RV remodeling manifests with a distinct molecular signature as determined by both transcriptomics and proteomics analyses. Our bioinformatics approach shows the Lysosome and Metabolism of Lipids pathways are associated with RVEF, the Dilated Cardiomyopathy and Arrhythmogenic Right Ventricular Cardiomyopathy pathways are mostly suppressed as RV hypertrophy progresses, and the Metabolism, Respiratory Electron Transport, and Citric Acid Cycle and Respiratory Electron Transport pathways are closely related to pathological RV dilation. Finally, a species comparison shows pigs more accurately recapitulate the human RVF metabolic molecular phenotype than rats. In summary, our multi-species cohort study nominates potential therapeutic targets for RVF, and raises the possibility that a porcine model could be used to test RV-directed therapeutics as a way to potentially improve the likelihood of successful translation to humans.

When defining molecules associated with RV function, we show NPC1 transcript and protein levels are negatively correlated with RVEF. NPC1 mutations cause NPC1 disease, a neurodegenerative and liver disorder characterized by cholesterol accumulation due to impairments in autophagic flux(26). Perhaps the dysregulation in NPC1 in PAB pigs promotes RVF due to alteration of autophagy. Interestingly, disruptions in autophagy are observed in multiple forms of left heart failure, and autophagy activation may have therapeutic effects (27,28). In dilated cardiomyopathy patients, increased abundance of autophagic vacuoles is a predictor of left ventricular reverse remodeling following medical therapy (29). However, modulation of autophagy in the setting of cardiac dysfunction needs to be considered carefully as some approaches may have detrimental effects. Induction of autophagy via AAV9-mediated overexpression of transcription factor EB (TFEB) enhances beneficial reverse cardiac remodeling after aortic debanding in the setting of left anterior descending (LAD) artery ligation. However, AAV9-TFEB accelerates heart failure progression and increases mortality if LAD ligation/aortic constriction mice are not debanded (30). Thus, the nature and duration of the physiological insult may be important for determining how and when to target autophagy to augment RV function.

The ETC is composed of five individual complexes that work together to facilitate oxidative phosphorylation(31). To provide a more thorough analysis of the ETC, we evaluated the transcriptomic and proteomic regulation of each complex and compared the results from rats and pigs as human samples did not identify as many ETC subunits. In the proteomics analysis, MCT rats have decreased abundance of all five ETC complexes (**Supplemental Figures 14-18**), which the transcriptomic analysis recapitulates. PAB pigs display mostly increased protein abundance of all five ETC complex pathways in proteomics analysis (**Supplemental Figures 14-18**). However, the transcriptomic analysis does not reveal uniform changes across all five ETC complexes as the abundance of transcripts in complexes II and V decrease, complex IV transcripts increase, and there is not a consistent pattern for transcripts in complexes I and II. These data suggest there are more profound impairments in ETC regulation in rodents than what is observed in pigs.

Ineffective FAO promotes RVF due to the critical role of FAO in ATP generation(32), but suppressed FAO may also contribute to RVF via lipotoxicity (23). Certainly, our data and others’ suggest there is disruption of FAO in rodents, pigs, and humans with RVF(33), but the directionality of FAO enzyme regulation seems to differ in these three species (**Figure 6**). Interestingly, there appears to be a conserved lipotoxic response in rodents and humans with RVF. In particular, there is accumulation of ceramides, a toxic lipid species that promotes cardiac dysfunction(32), in both rodent and human RVF (10,23,34). Perhaps suppression of lipotoxicity may be an added benefit of augmenting FAO in the setting of RVF. However, further work is needed to understand the interplay between impaired FAO and lipotoxicity in RVF, especially in more translational large animal models.

While our data suggests FAO is more robustly suppressed in rodent as compared to porcine and human RVF, there are pharmacological approaches, lifestyle interventions, and supplementation studies that suggest either suppression or activation of FAO imparts RV-enhancing effects in both rodents and humans(35). FAO is increased in rodents following pulmonary artery banding, and inhibition of FAO with ranolazine and trimetazidine augments RV function(36). This finding is conserved in humans as a small clinical trial shows ranolazine increases RV ejection fraction in pulmonary hypertension-mediated RVF(37). On the other hand, activation of AMP-kinase, a master regulator of mitochondrial FAO, via small molecule With-No-Lysine Kinase inhibition or intermittent fasting normalizes FAO protein abundance and improves RV performance in MCT and Sugen-hypoxia rats(10,34,38). In both a genetic mouse model of PAH and PAB mice, L-carnitine treatment ameliorates RV lipid accumulation and RV dysfunction(39), again highlighting the beneficial effects of FAO activation. Moreover, the use of the peroxisome proliferator activated receptor gamma agonist, pioglitazone, restores FAO protein regulation and improves RV function in Sugen-hypoxia rats, but that effect may be partially explained by a significant reduction in pulmonary vascular disease(40). Finally, a phase II clinical trial shows metformin, a FAO activator, increases RV fractional area change in PAH-mediated RV dysfunction(41). Certainly, there are beneficial RV effects with both FAO activation and inhibition in rodents and humans. Perhaps the degree of FAO dysregulation, whether induced or suppressed, will determine which approach will have the greatest efficacy for RVF.

Although there are many divergent molecular responses when comparing the three species, alterations in the Dilated Cardiomyopathy and Arrhythmogenic RV Cardiomyopathy (ARVC) pathways appear conserved as proteins in these two pathways are predominately downregulated in all three species (**Supplemental Figures 19-20**). This suggests that Dilated Cardiomyopathy and ARVC proteins may be important targets when considering RV therapeutics, and there are data to support this hypothesis. First, our and Boucherat *et al*., proteomics data show junctophilin-2, an ARVC protein that maintains t-tubule structure and proper excitation-contraction coupling(42), levels are lowered in both PAB pigs and humans with RVF(13,42,43). Combatting junctophilin-2 downregulation improves RV function in rodent RVF(10,13,43), further supporting its key role in maintaining RV contractility. Second, the ARVC pathway is enriched in desmosomal proteins, and mutations in these proteins cause a RV-predominant cardiomyopathy(44). Our previous proteomics analysis shows numerous desmosomal proteins are reduced in the RV of MCT rats(13), which provides additional support that desmosomal proteins play a key role in the RV. Finally, there is mislocalization of the desmosomal/gap junction protein connexin-43 in both rodent and large animal RVF (45,46), again demonstrating a conserved response across species. In summary, the consistent finding of dysregulation of ARVC and Dilated Cardiomyopathy pathway proteins highlights their potential as therapeutic targets for RVF.

In summary, we show PAB induces RV remodeling and dysfunction in pigs. We demonstrate Lysosomal, Lipid Metabolism, FAO, and ETC pathways are associated with pathogenic RV remodeling and impaired RV function. Our interspecies comparison suggests the Dilated Cardiomyopathy and ARVC pathways are comparably dysregulated in rats, pigs, and humans, but the porcine model is more similar to humans with specific regards to metabolic/mitochondrial regulation than rodents. Thus, the porcine model may serve an important role in evaluating RV-directed therapies, once lead compounds or approaches are identified in rodents, to hopefully increase the likelihood of translation to humans.

### Limitations

Our manuscript has important limitations that we must acknowledge. The PAB model in piglets caused moderate RV dysfunction whereas the rat MCT model results in significant RV dysfunction and death. The divergent changes in protein abundances within specific pathways may reflect variation in disease severity rather than simply species differences. While we only evaluated the transcriptomic and proteomic changes in the MCT model, the MCT and Sugen-hypoxia models have highly similar transcriptomic signatures(47). Moreover, these two rat models exhibit conserved changes in ETC and FAO proteins in the RV(38). All animal models only used male sex, but the human data included females. However, the impact of biological sex on the molecular mechanisms of RVF in large animal models still needs to be investigated.

### Clinical Perspectives

Competency in Medical Knowledge: RVF is a risk factor for death in multiple cardiovascular diseases, yet there are currently no RV-specific therapies. Here, we show our porcine model recapitulates many molecular phenotypes of human RVF, suggesting it has utility for understanding mechanisms of human RVF.

Translational Outlook: We identified multiple pathways associated with adverse RV remodeling and RV dysfunction. Future studies evaluating therapeutics targeting these important molecular mechanisms may allow for the development of RVF combatting pharmaceuticals.

---

**Supplemental Figure 1:**
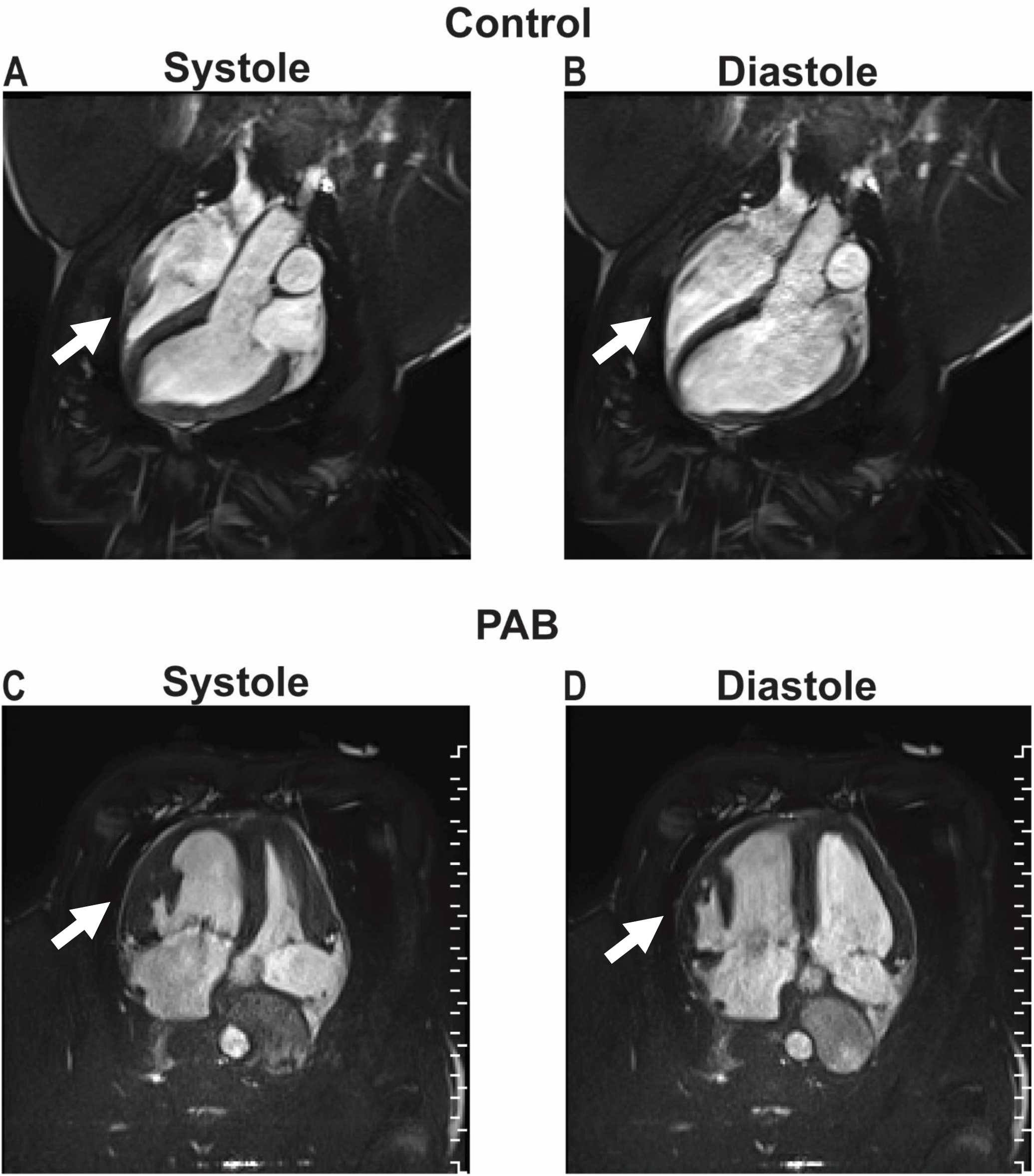
Representative still images from cMRI of control (A)/(B) and PAB (C)/(D) piglets at end-systole (A)/(C), and end-diastole (B)/(D). Arrows denote the right ventricle in each of the panels.

**Supplemental Figure 2:**
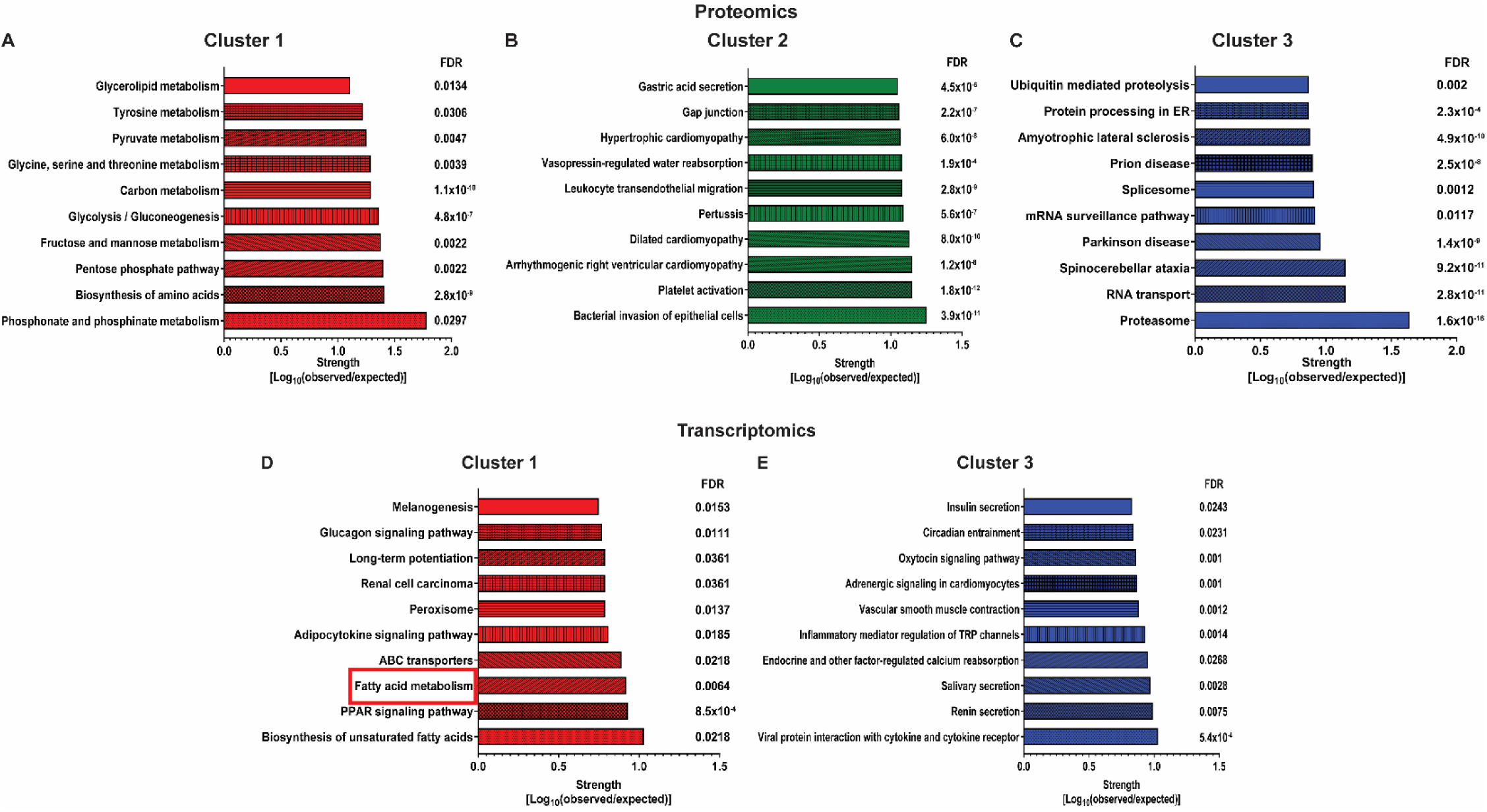
KEGG pathways identified from proteins (A), (B), (C) and transcripts (D), (E) positively correlated with RVEF. The Fatty Acid Metabolism pathway was associated with RVEF in both the proteomic and transcriptomic analyses.

**Supplemental Figure 3:**
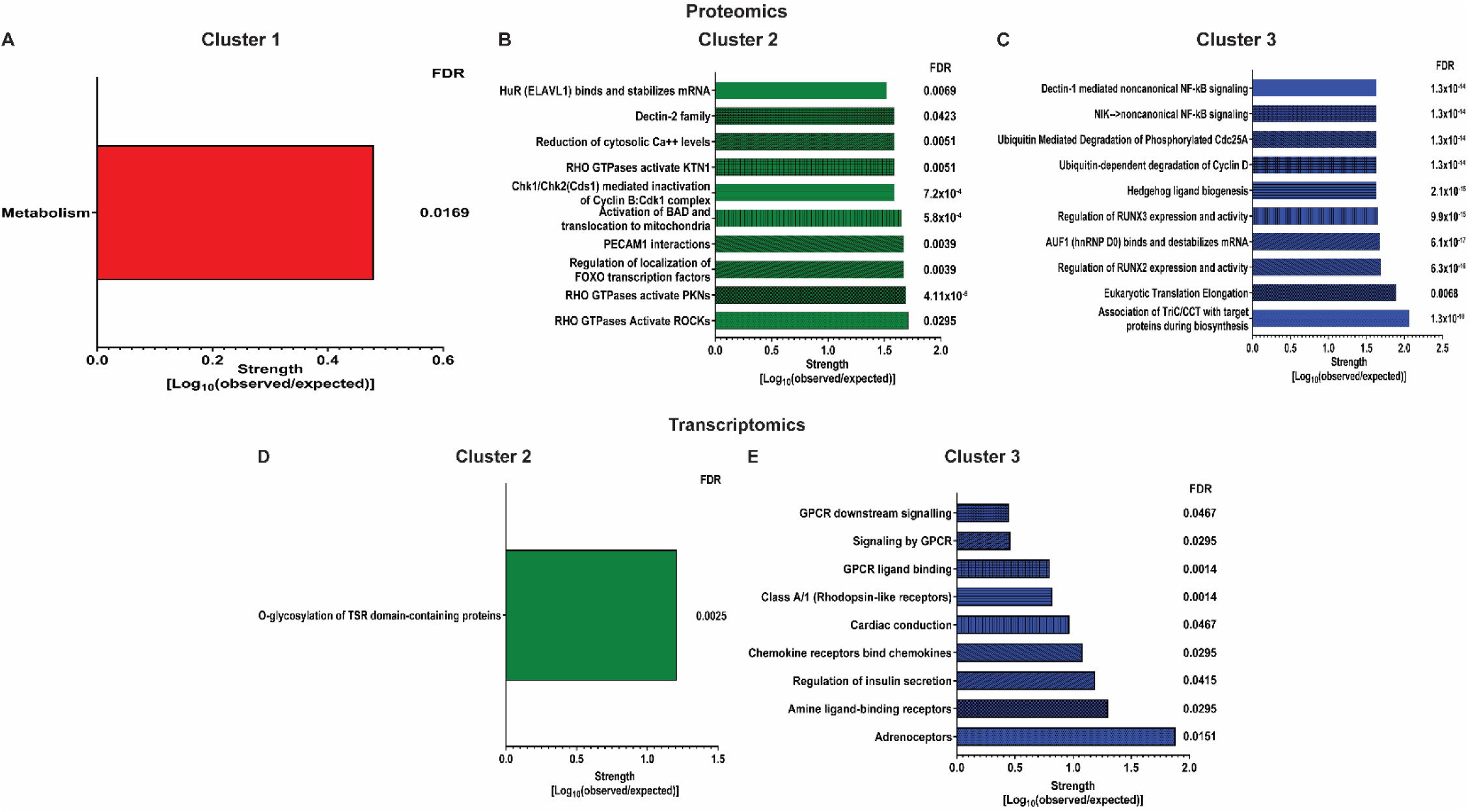
Reactome pathways identified from proteins (A), (B), (C) and transcripts (D), (E) positively correlated with RVEF.

**Supplemental Figure 4:**
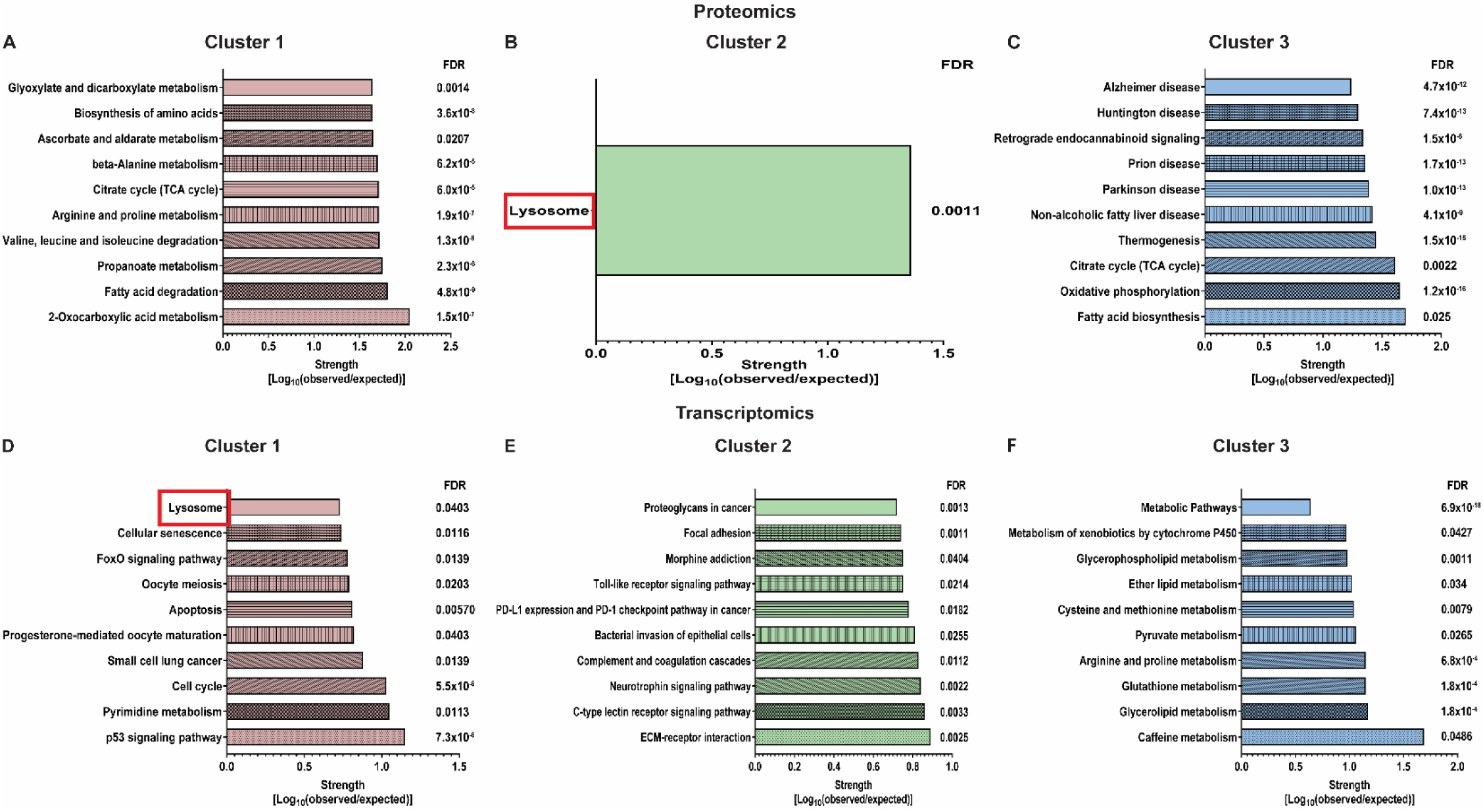
KEGG pathways identified from proteins (A), (B), (C) and transcripts (D), (E), (F) negatively correlated with RVEF. The Lysosome pathway was identified in both the proteomics and transcriptomics analyses.

**Supplemental Figure 5:**
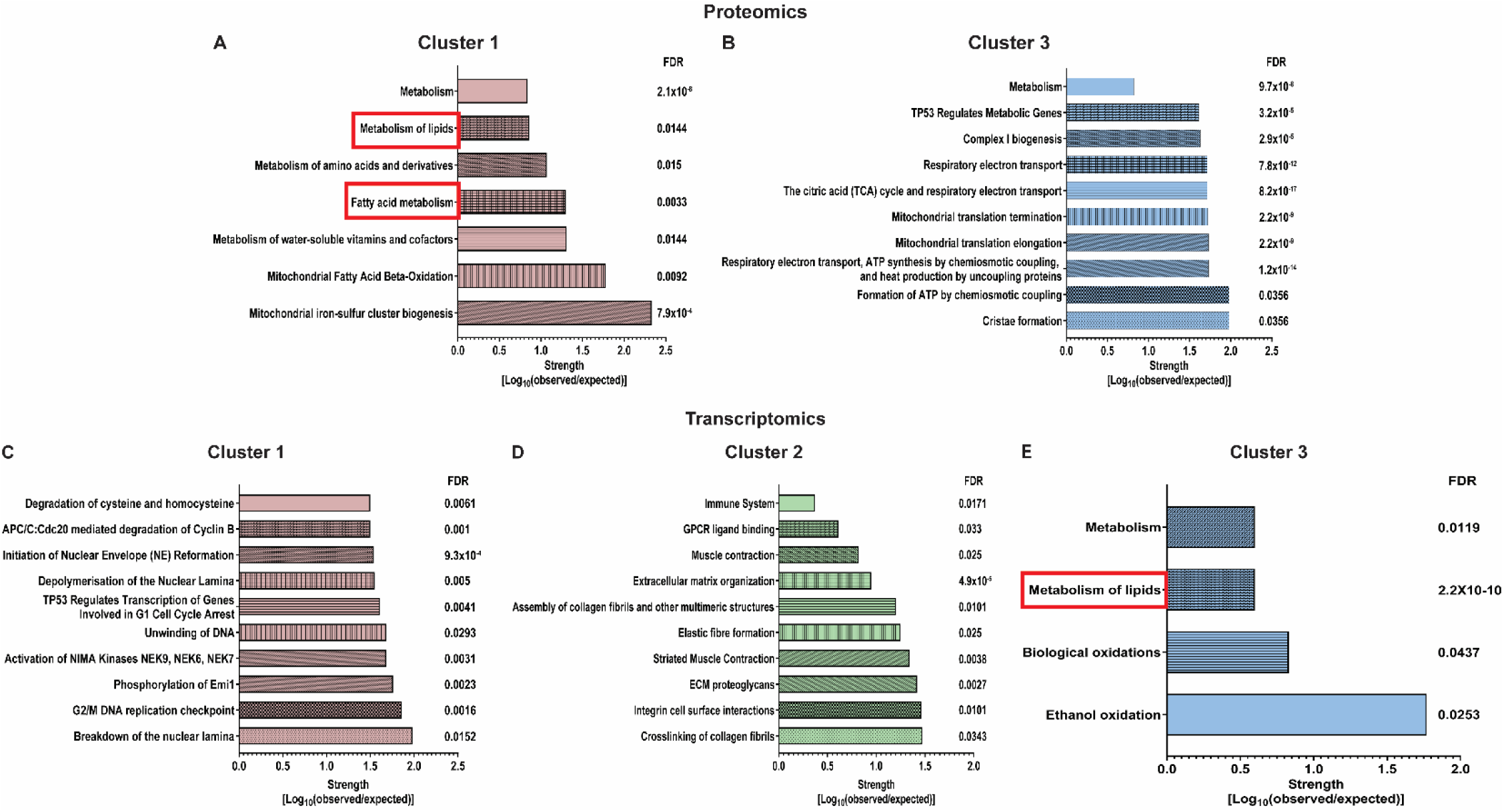
Reactome pathways identified from proteins (A), (B) and transcripts (C), (D), (E) negatively correlated with RVEF. The Metabolism of Lipids pathway was identified in both the proteomics and transcriptomics analyses. The Fatty Acid Metabolism pathway was associated with RVEF in both the proteomics and transcriptomics analyses.

**Supplemental Figure 6:**
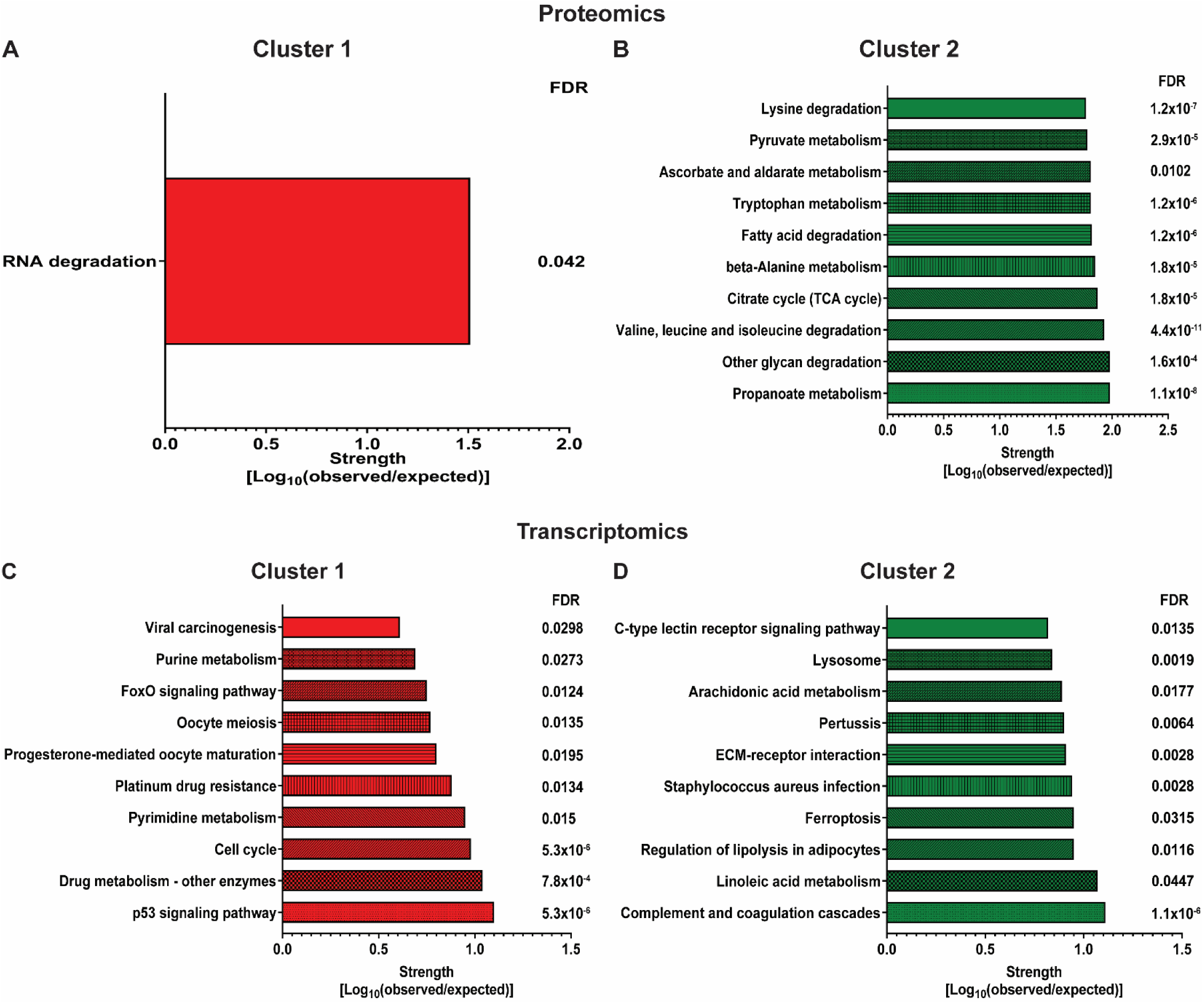
KEGG pathways identified from proteins (A), (B), and transcripts (C), (D) positively correlated with RV mass.

**Supplemental Figure 7:**
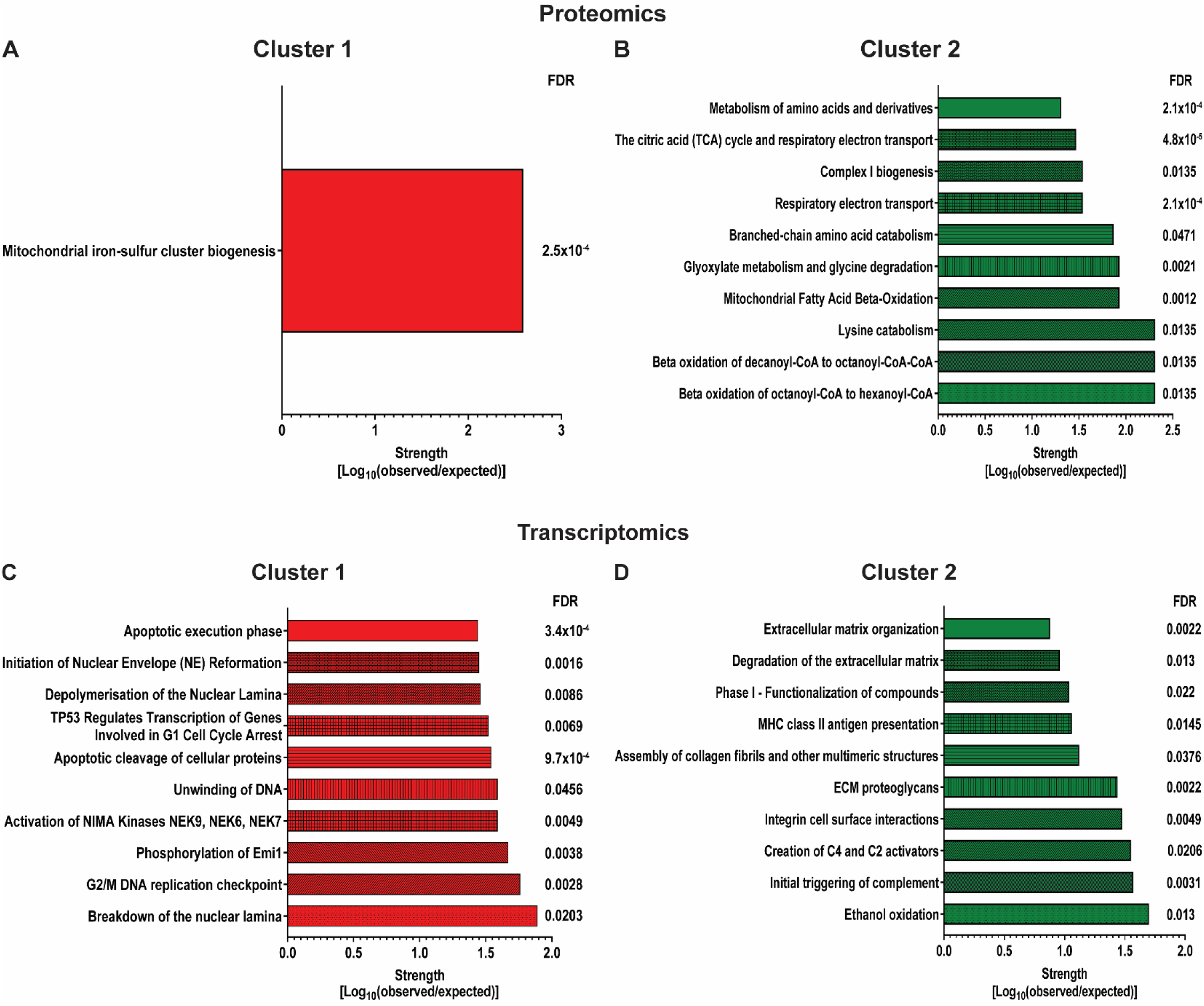
Reactome pathways identified from proteins (A), (B), and transcripts (C), (D) positively correlated with RV mass.

**Supplemental Figure 8:**
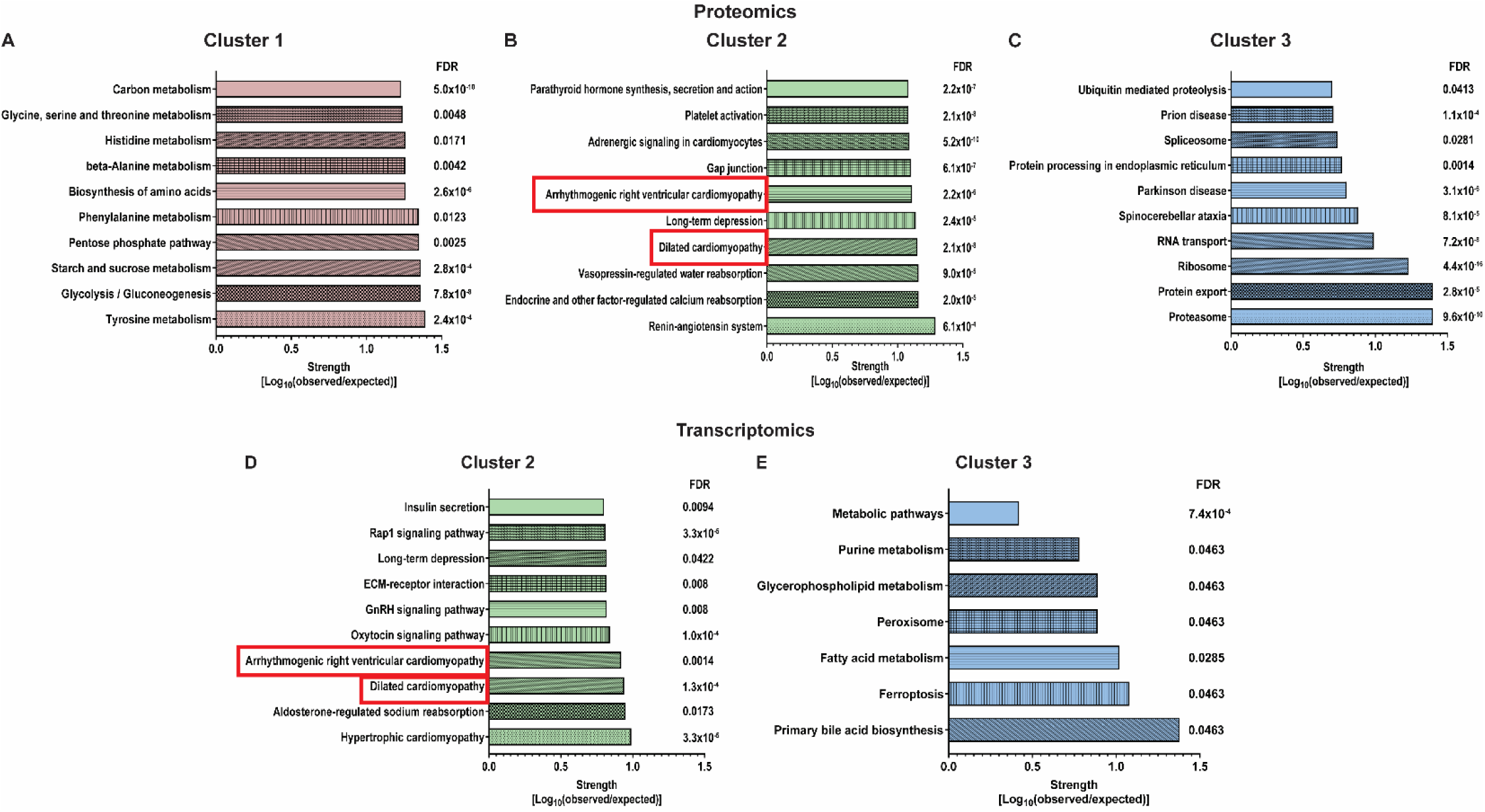
KEGG pathways identified from proteins (A), (B), (C), and transcripts (D), (E) negatively correlated with RV mass. The Dilated Cardiomyopathy and Arrhythmogenic Right Ventricular Cardiomyopathy pathways were identified in both the proteomics and transcriptomics analysis.

**Supplemental Figure 9:**
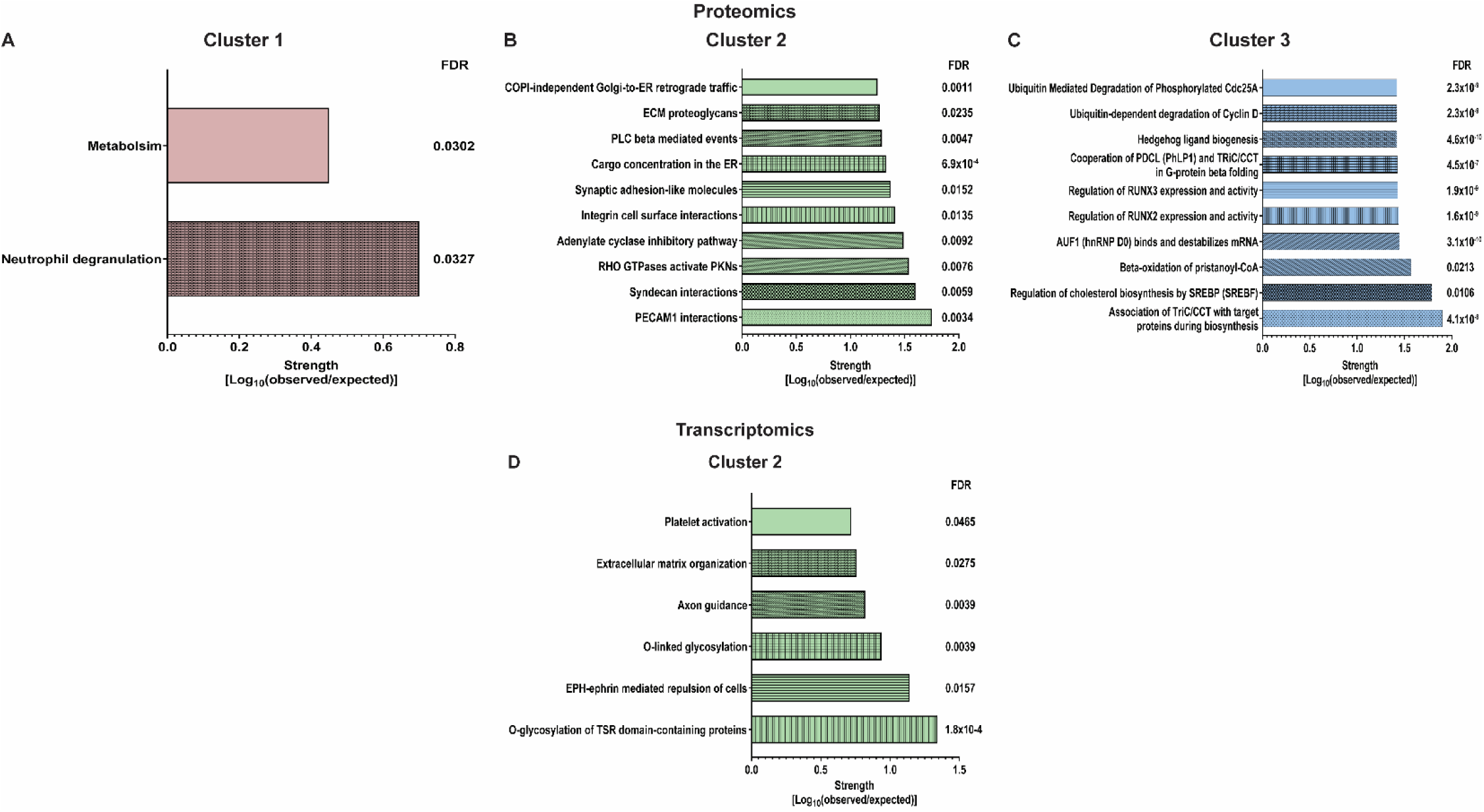
Reactome pathways identified from proteins (A), (B), (C) and transcripts (D) negatively correlated with RV mass.

**Supplemental Figure 10:**
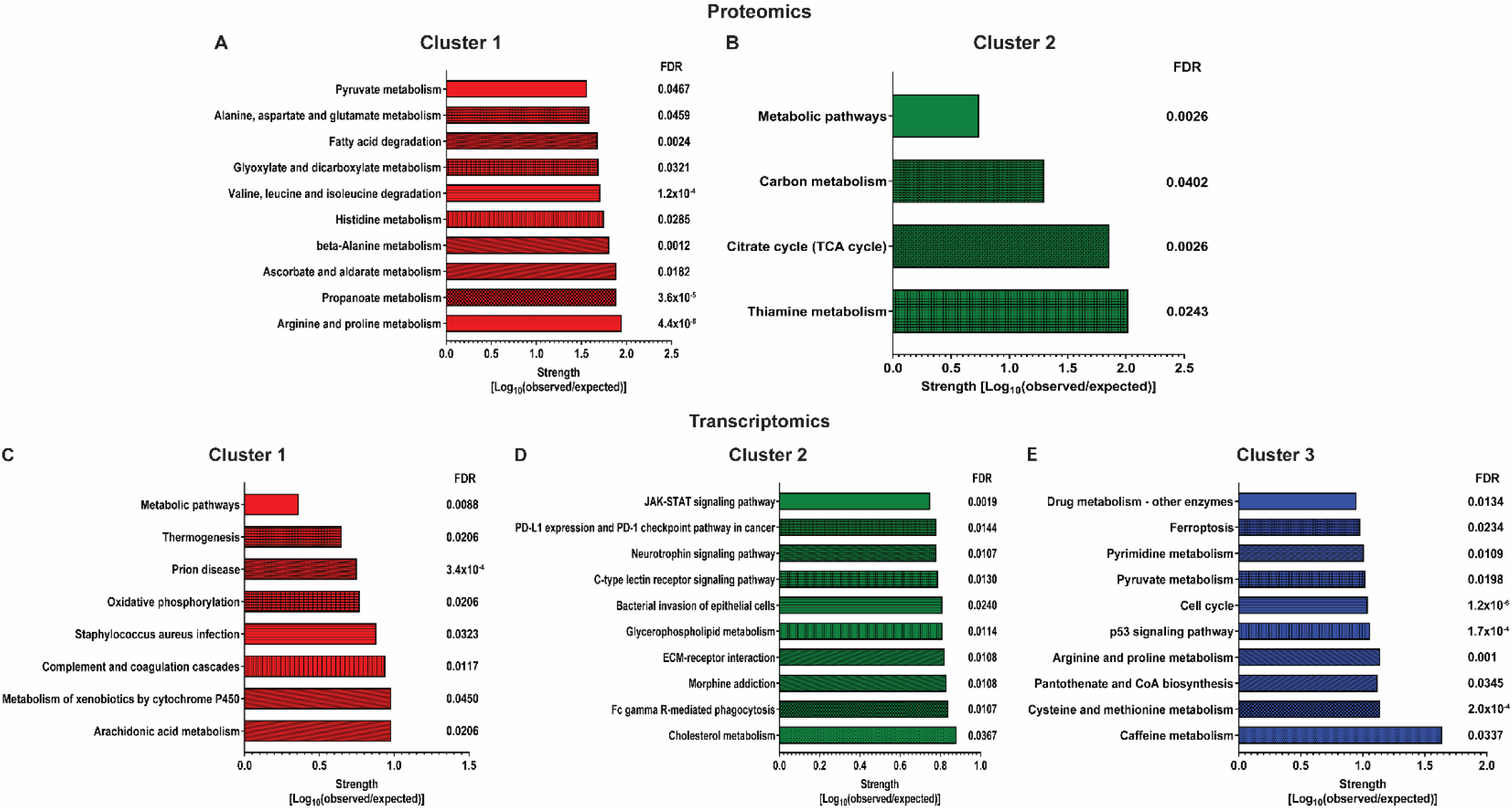
KEGG pathways identified from proteins (A), (B) and transcripts (C), (D), (E) positively correlated with RV ESV.

**Supplemental Figure 11:**
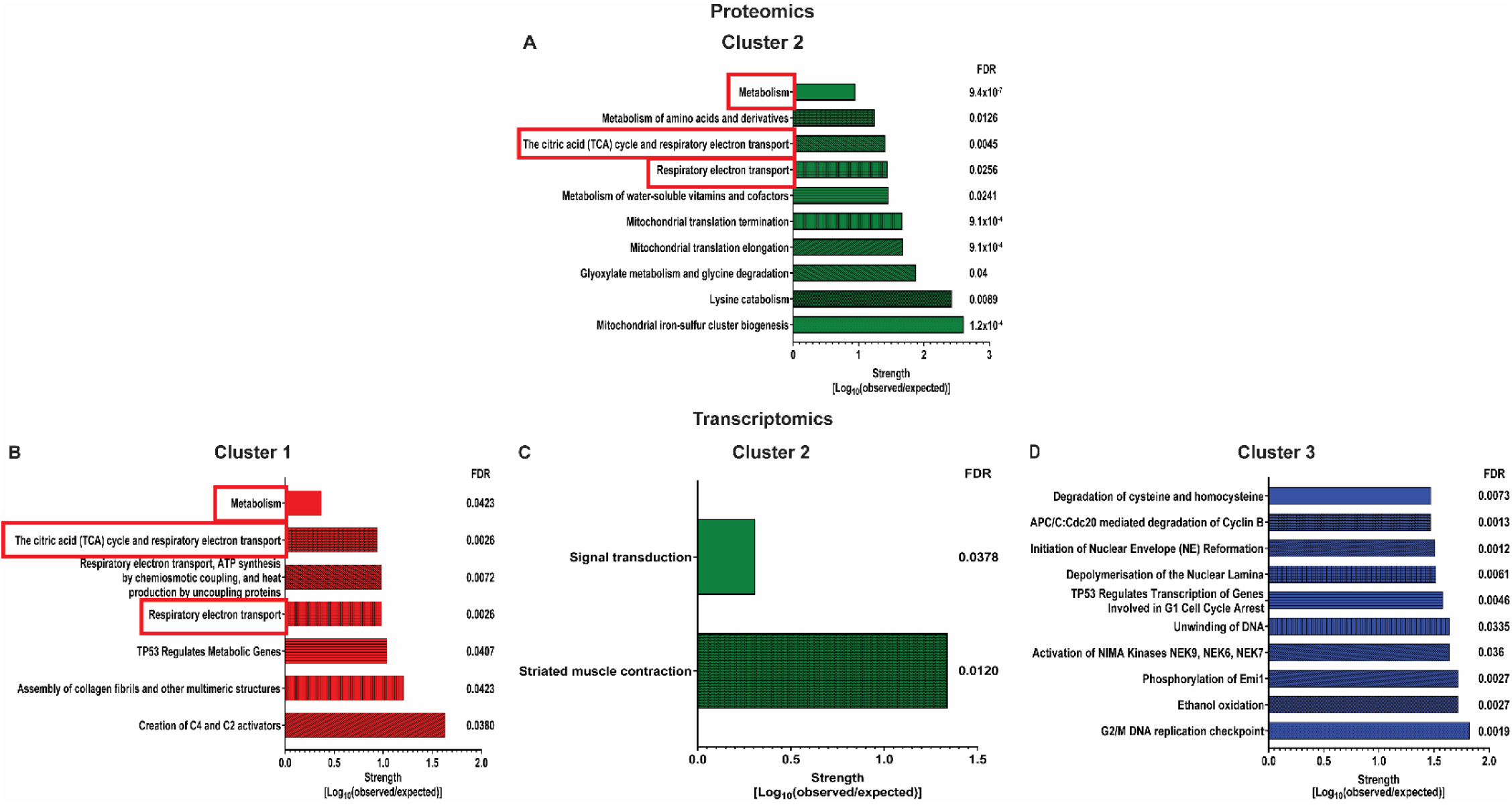
Reactome pathways identified from proteins (A) and transcripts (B), (C), (D) positively correlated with RV ESV. The Citric Acid (TCA Cycle) and Respiratory Electron Transport, Metabolism, and Respiratory Electron Transport pathways were identified in both the proteomics and transcriptomics analyses.

**Supplemental Figure 12:**
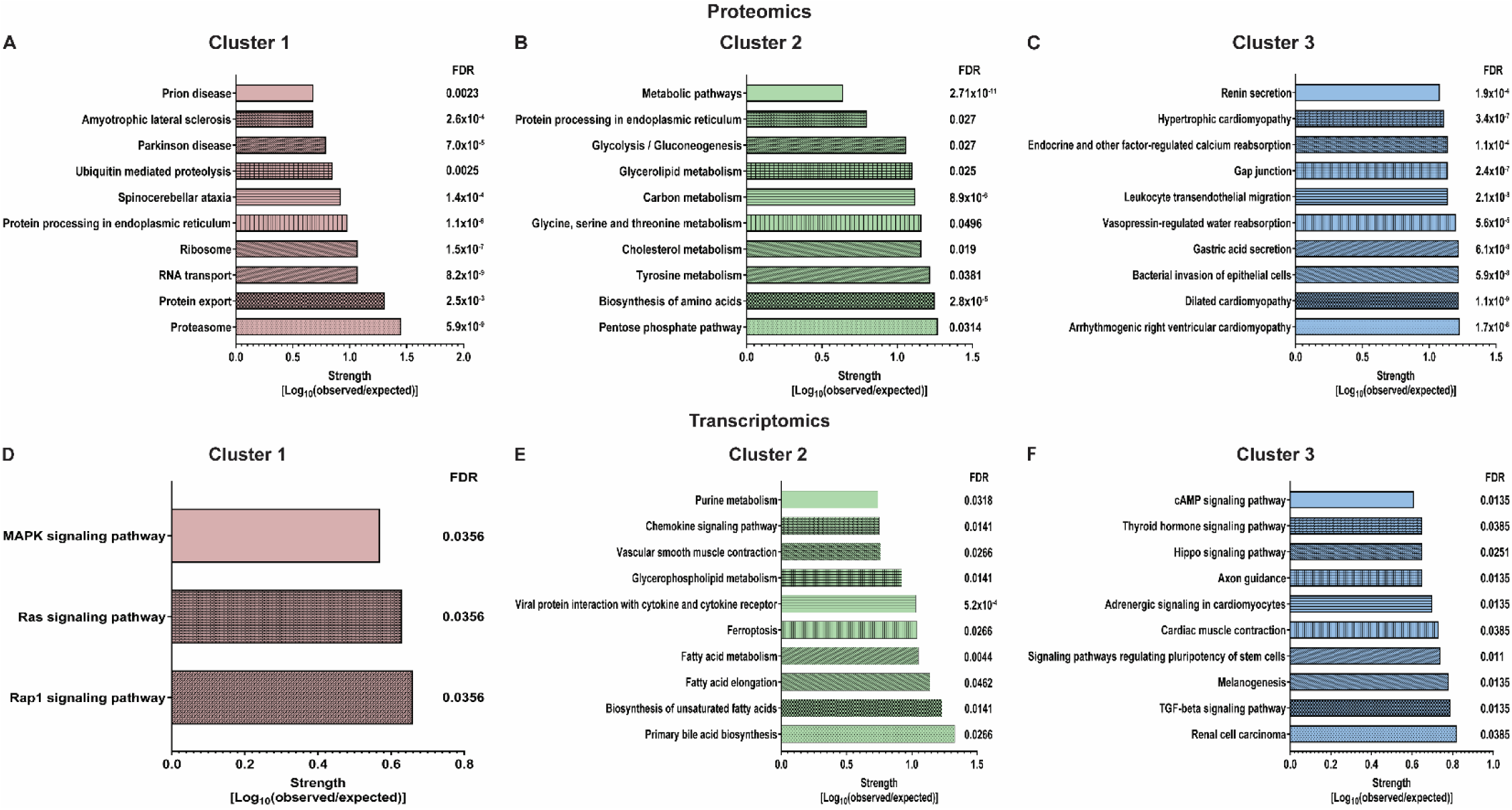
KEGG pathways identified from proteins (A), (B), (C) and transcripts (D), (E), (F) negatively correlated with RV ESV.

**Supplemental Figure 13:**
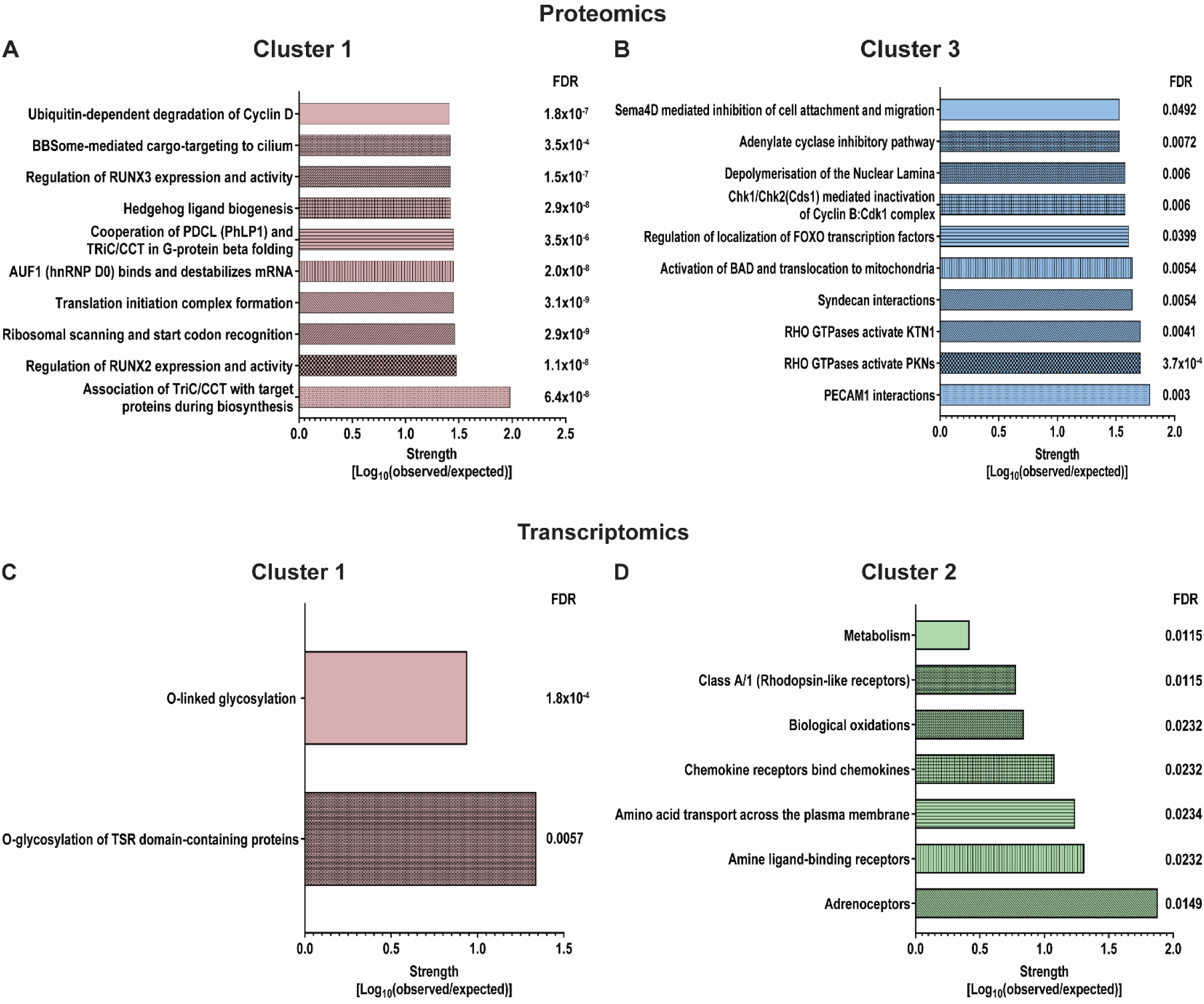
Reactome pathways identified from proteins (A), (B) and transcripts (C), (D) negatively correlated with RV ESV.

**Supplemental Figure 14:**
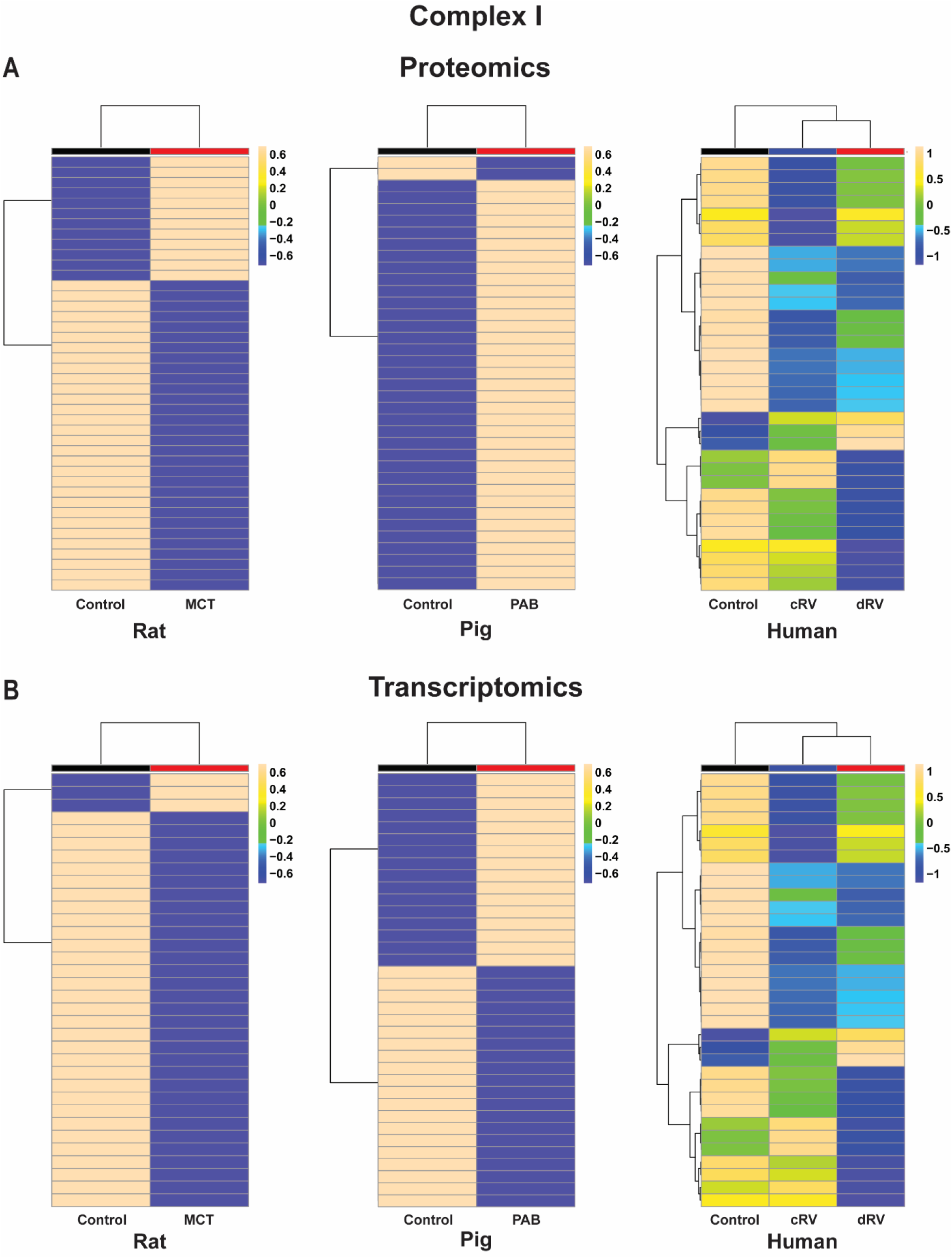
Hierarchical cluster analyses of (**A**) proteomics and (**B**) transcriptomics data from rat (left), pig (center) and human (right) demonstrate inter-species differences in the expression of complex I of the electron transport chain.

**Supplemental Figure 15:**
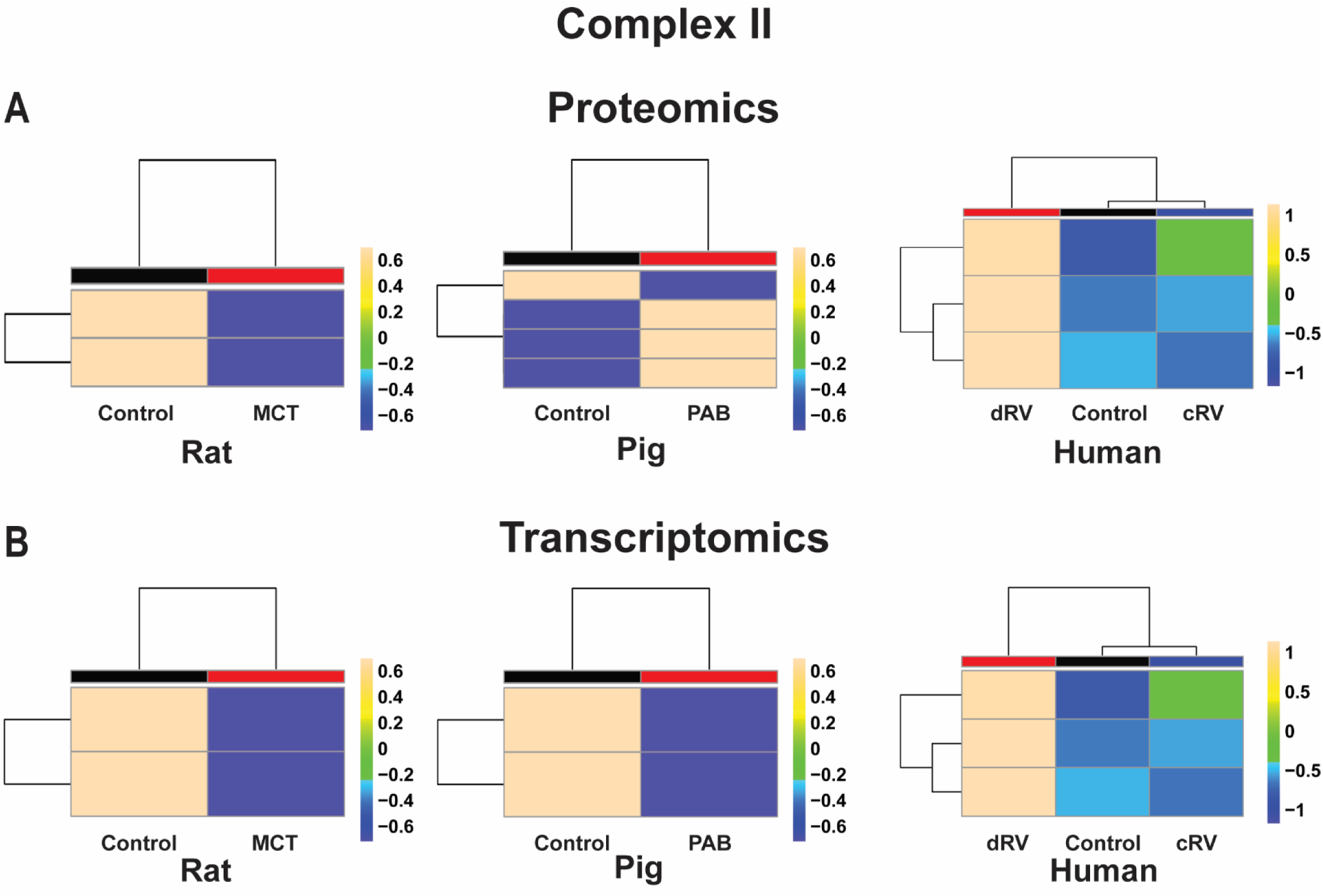
Hierarchical cluster analyses of (**A**) proteomics and (**B**) transcriptomics data from rat (left), pig (center) and human (right) demonstrate inter-species differences in the expression of complex II of the electron transport chain.

**Supplemental Figure 16:**
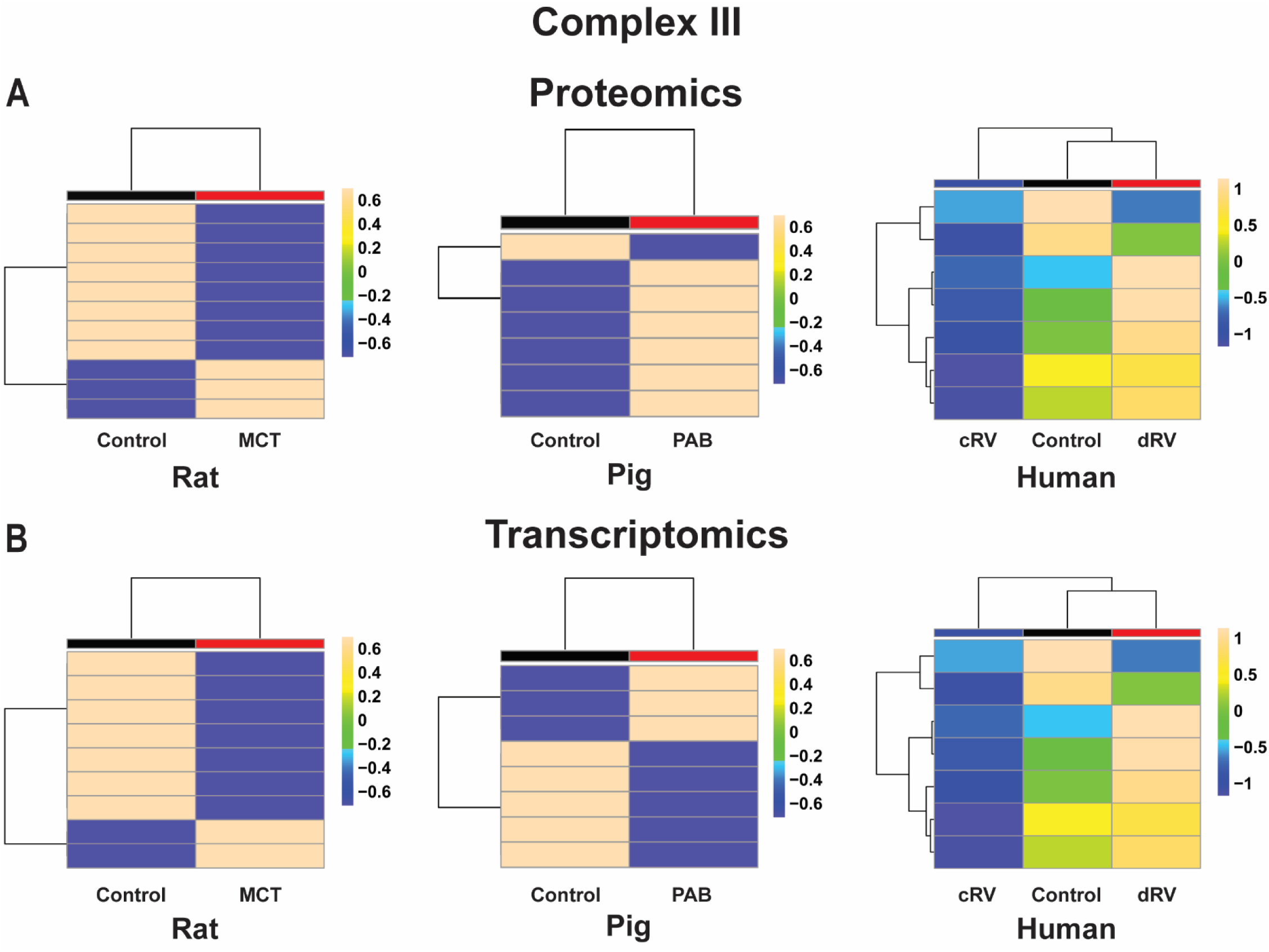
Hierarchical cluster analyses of (**A**) proteomics and (**B**) transcriptomics data from rat (left), pig (center) and human (right) demonstrate inter-species differences in the expression of complex III of the electron transport chain.

**Supplemental Figure 17:**
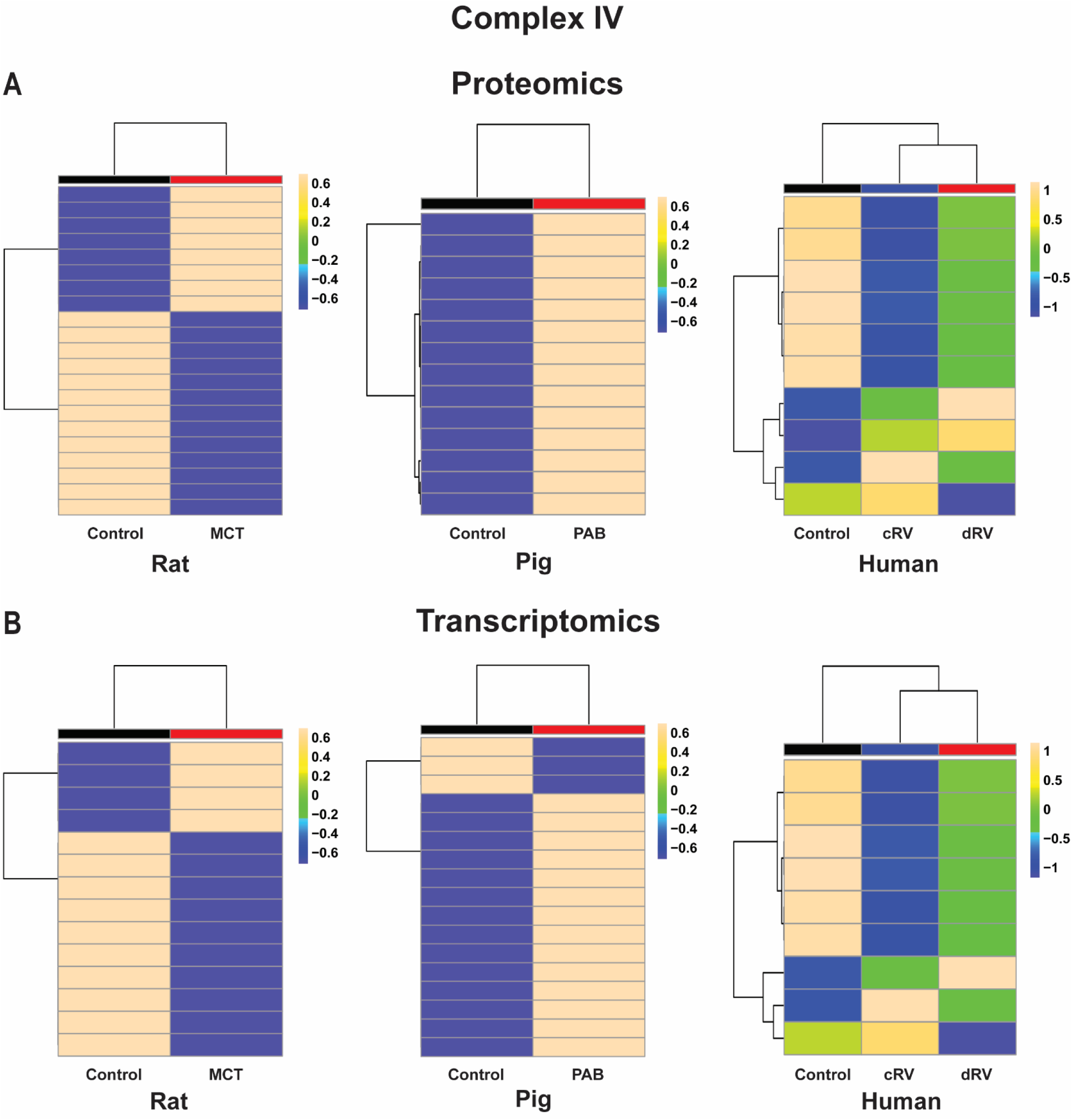
Hierarchical cluster analyses of (**A**) proteomics and (**B**) transcriptomics data from rat (left), pig (center) and human (right) demonstrate inter-species differences in the expression of complex IV of the electron transport chain.

**Supplemental Figure 18:**
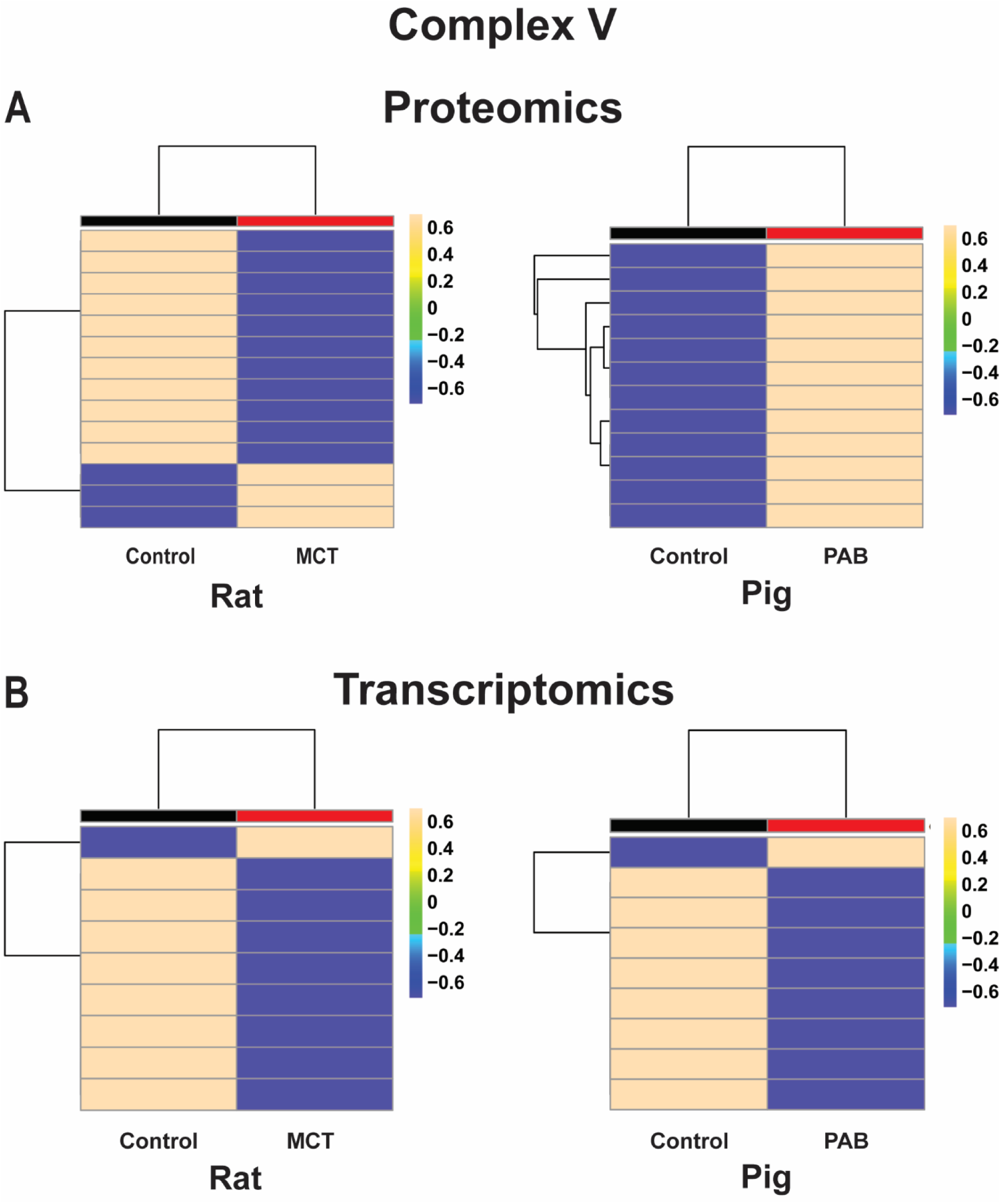
Hierarchical cluster analyses of (**A**) proteomics and (**B**) transcriptomics data from rat (left), pig (right) demonstrate inter-species differences in the expression of complex V of the electron transport chain. The components of complex V were not measured in the human data set.

**Supplemental Figure 19:**
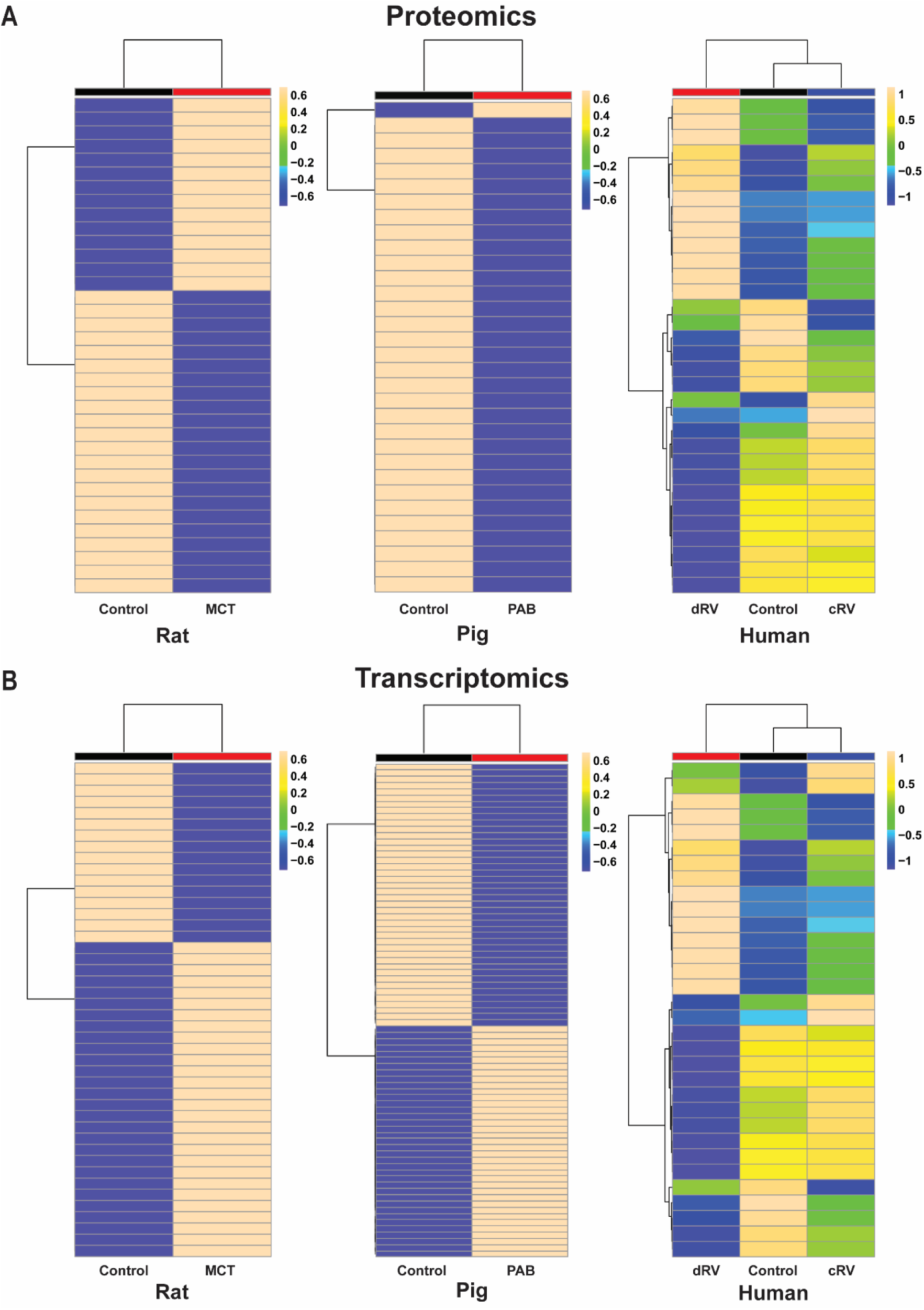
Hierarchical cluster analyses of (**A**) proteomics and (**B**) transcriptomics data from rat (left), pig (center) and human (right) demonstrate inter-species similarities in the expression of the Dilated Cardiomyopathy pathway.

**Supplemental Figure 20:**
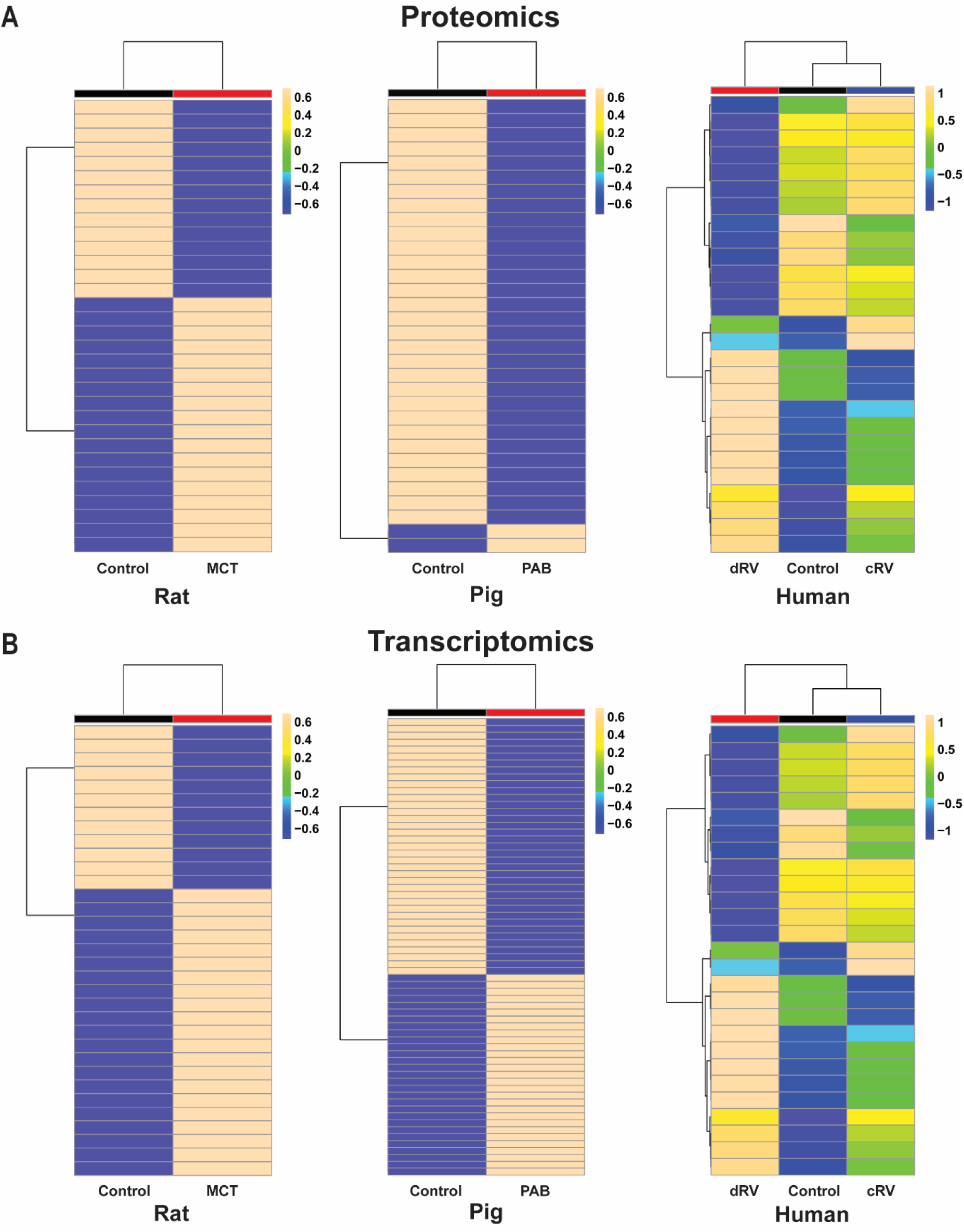
Hierarchical cluster analyses of (**A**) proteomics and (**B**) transcriptomics data from rat (left), pig (center) and human (right) demonstrate inter-species similarities in the expression of the Arrhythmogenic Right Ventricular Cardiomyopathy pathway.

## References

1. Ghio S, Gavazzi A, Campana C et al. Independent and additive prognostic value of right ventricular systolic function and pulmonary artery pressure in patients with chronic heart failure. J Am Coll Cardiol 2001;37:183–8.

2. Tadic M, Nita N, Schneider L et al. The Predictive Value of Right Ventricular Longitudinal Strain in Pulmonary Hypertension, Heart Failure, and Valvular Diseases. Front Cardiovasc Med 2021;8:698158.

3. Friedberg MK, Reddy S. Right ventricular failure in congenital heart disease. Curr Opin Pediatr 2019;31:604–610.

4. Cho YK, Ma JS. Right ventricular failure in congenital heart disease. Korean J Pediatr 2013;56:101–6.

5. Vonk Noordegraaf A, Westerhof BE, Westerhof N. The Relationship Between the Right Ventricle and its Load in Pulmonary Hypertension. J Am Coll Cardiol 2017;69:236–243.

6. Voelkel NF, Gomez-Arroyo J, Abbate A, Bogaard HJ, Nicolls MR. Pathobiology of pulmonary arterial hypertension and right ventricular failure. Eur Respir J 2012;40:1555–65.

7. Prisco SZ, Thenappan T, Prins KW. Treatment Targets for Right Ventricular Dysfunction in Pulmonary Arterial Hypertension. JACC Basic Transl Sci 2020;5:1244–1260.

8. Lunney JK, Van Goor A, Walker KE, Hailstock T, Franklin J, Dai C. Importance of the pig as a human biomedical model. Sci Transl Med 2021;13:eabd5758.

9. Provencher S, Archer SL, Ramirez FD et al. Standards and Methodological Rigor in Pulmonary Arterial Hypertension Preclinical and Translational Research. Circ Res 2018;122:1021–1032.

10. Prisco SZ, Eklund M, Raveendran R, Thenappan T, Prins KW. With No Lysine Kinase 1 Promotes Metabolic Derangements and RV Dysfunction in Pulmonary Arterial Hypertension. JACC Basic Transl Sci 2021;6:834–850.

11. Boucherat O, Yokokawa T, Krishna V et al. Identification of LTBP-2 as a plasma biomarker for right ventricular dysfunction in human pulmonary arterial hypertension. Nature Cardiovascular Research 2022;1:748–760.

12. Azakie A, Carney JP, Lahti MT et al. Porcine Model of the Arterial Switch Operation: Implications for Unique Strategies in the Management of Hypoplastic Left Ventricles. Pediatr Cardiol 2021;42:501–509.

13. Prisco SZ, Hartweck LM, Rose L et al. Inflammatory Glycoprotein 130 Signaling Links Changes in Microtubules and Junctophilin-2 to Altered Mitochondrial Metabolism and Right Ventricular Contractility. Circ Heart Fail 2022;15:e008574.

14. Baller J, Kono T, Herman A, Zhang Y. CHURP: A Lightweight CLI Framework to Enable Novice Users to Analyze Sequencing Datasets in Parallel. In Proceedings of the Practice and Experience in Advanced Research Computing on Rise of the Machines (learning). (PEARC ’19) Association for Computing Machinery. New York, NY, USA, 2019:1–5.

15. Kim D, Paggi JM, Park C, Bennett C, Salzberg SL. Graph-based genome alignment and genotyping with HISAT2 and HISAT-genotype. Nat Biotechnol 2019;37:907–915.

16. Love MI, Huber W, Anders S. Moderated estimation of fold change and dispersion for RNA-seq data with DESeq2. Genome Biol 2014;15:550.

17. Kolde R. Pheatmap: pretty heatmaps. R package version 2012;1:726.

18. Blighe K, Rana S, Lewis M. EnhancedVolcano: publication-ready volcano plots with enhanced colouring and labeling. R Package version 1.14.0 ed, 2022.

19. Chong J, Yamamoto M, Xia J. MetaboAnalystR 2.0: From Raw Spectra to Biological Insights. Metabolites 2019;9.

20. Szklarczyk D, Franceschini A, Wyder S et al. STRING v10: protein-protein interaction networks, integrated over the tree of life. Nucleic Acids Res 2015;43:D447–52.

21. Tello K, Dalmer A, Axmann J et al. Reserve of Right Ventricular-Arterial Coupling in the Setting of Chronic Overload. Circ Heart Fail 2019;12:e005512.

22. Kwon HJ, Abi-Mosleh L, Wang ML et al. Structure of N-terminal domain of NPC1 reveals distinct subdomains for binding and transfer of cholesterol. Cell 2009;137:1213–24.

23. Brittain EL, Talati M, Fessel JP et al. Fatty Acid Metabolic Defects and Right Ventricular Lipotoxicity in Human Pulmonary Arterial Hypertension. Circulation 2016;133:1936–44.

24. Flam E, Jang C, Murashige D et al. Integrated landscape of cardiac metabolism in end-stage human nonischemic dilated cardiomyopathy. Nature Cardiovascular Research, 2022:817–829.

25. Hindmarch CCT, Tian L, Xiong PY et al. An integrated proteomic and transcriptomic signature of the failing right ventricle in monocrotaline induced pulmonary arterial hypertension in male rats. Front Physiol 2022;13:966454.

26. Sarkar S, Carroll B, Buganim Y et al. Impaired autophagy in the lipid-storage disorder Niemann-Pick type C1 disease. Cell Rep 2013;5:1302–15.

27. Schiattarella GG, Hill JA. Therapeutic targeting of autophagy in cardiovascular disease. J Mol Cell Cardiol 2016;95:86–93.

28. Choi AM, Ryter SW, Levine B. Autophagy in human health and disease. N Engl J Med 2013;368:651–62.

29. Kanamori H, Yoshida A, Naruse G et al. Impact of Autophagy on Prognosis of Patients With Dilated Cardiomyopathy. J Am Coll Cardiol 2022;79:789–801.

30. Evans S, Ma X, Wang X et al. Targeting the Autophagy-Lysosome Pathway in a Pathophysiologically Relevant Murine Model of Reversible Heart Failure. JACC: Basic to Translational Science;0.

31. Acin-Perez R, Benador IY, Petcherski A et al. A novel approach to measure mitochondrial respiration in frozen biological samples. EMBO J 2020;39:e104073.

32. Lopaschuk GD, Karwi QG, Tian R, Wende AR, Abel ED. Cardiac Energy Metabolism in Heart Failure. Circ Res 2021;128:1487–1513.

33. Agrawal V, Lahm T, Hansmann G, Hemnes AR. Molecular mechanisms of right ventricular dysfunction in pulmonary arterial hypertension: focus on the coronary vasculature, sex hormones, and glucose/lipid metabolism. Cardiovasc Diagn Ther 2020;10:1522–1540.

34. Prisco SZ, Eklund M, Moutsoglou DM et al. Intermittent Fasting Enhances Right Ventricular Function in Preclinical Pulmonary Arterial Hypertension. J Am Heart Assoc 2021;10:e022722.

35. Talati M, Hemnes A. Fatty acid metabolism in pulmonary arterial hypertension: role in right ventricular dysfunction and hypertrophy. Pulm Circ 2015;5:269–78.

36. Fang YH, Piao L, Hong Z et al. Therapeutic inhibition of fatty acid oxidation in right ventricular hypertrophy: exploiting Randle’s cycle. J Mol Med (Berl) 2012;90:31–43.

37. Han Y, Forfia P, Vaidya A et al. Ranolazine Improves Right Ventricular Function in Patients With Precapillary Pulmonary Hypertension: Results From a Double-Blind, Randomized, Placebo-Controlled Trial. J Card Fail 2021;27:253–257.

38. Kazmirczak F, Hartweck LM, Vogel NT et al. Intermittent Fasting Activates AMP-Kinase to Restructure Right Ventricular Lipid Metabolism and Microtubules in Two Rodent Models of Pulmonary Arterial Hypertension. JACC: Basic to Translational Science 2022:2022.03.07.483333.

39. Agrawal V, Hemnes AR, Shelburne NJ et al. l-Carnitine therapy improves right heart dysfunction through Cpt1- dependent fatty acid oxidation. Pulm Circ 2022;12:e12107.

40. Legchenko E, Chouvarine P, Borchert P et al. PPARγ agonist pioglitazone reverses pulmonary hypertension and prevents right heart failure via fatty acid oxidation. Sci Transl Med 2018;10.

41. Brittain EL, Niswender K, Agrawal V et al. Mechanistic Phase II Clinical Trial of Metformin in Pulmonary Arterial Hypertension. J Am Heart Assoc 2020;9:e018349.

42. Zhang C, Chen B, Guo A et al. Microtubule-mediated defects in junctophilin-2 trafficking contribute to myocyte transverse-tubule remodeling and Ca2+ handling dysfunction in heart failure. Circulation 2014;129:1742–50.

43. Prins KW, Tian L, Wu D, Thenappan T, Metzger JM, Archer SL. Colchicine Depolymerizes Microtubules, Increases Junctophilin-2, and Improves Right Ventricular Function in Experimental Pulmonary Arterial Hypertension. J Am Heart Assoc 2017;6.

44. Yang Z, Bowles NE, Scherer SE et al. Desmosomal dysfunction due to mutations in desmoplakin causes arrhythmogenic right ventricular dysplasia/cardiomyopathy. Circ Res 2006;99:646–55.

45. Chkourko HS, Guerrero-Serna G, Lin X et al. Remodeling of mechanical junctions and of microtubule-associated proteins accompany cardiac connexin43 lateralization. Heart Rhythm 2012;9:1133–1140.e6.

46. Uzzaman M, Honjo H, Takagishi Y et al. Remodeling of gap junctional coupling in hypertrophied right ventricles of rats with monocrotaline-induced pulmonary hypertension. Circ Res 2000;86:871–8.

47. Park JF, Clark VR, Banerjee S et al. Transcriptomic Analysis of Right Ventricular Remodeling in Two Rat Models of Pulmonary Hypertension: Identification and Validation of Epithelial-to-Mesenchymal Transition in Human Right Ventricular Failure. Circ Heart Fail 2021;14:e007058.

